# Basement membrane regulates fibronectin organization using sliding focal adhesions driven by a contractile winch

**DOI:** 10.1101/618686

**Authors:** Jiaoyang Lu, Andrew D. Doyle, Yoshinari Shinsato, Shaohe Wang, Molly A. Bodendorfer, Minhua Zheng, Kenneth M. Yamada

## Abstract

We have discovered that basement membrane and its major components can induce rapid, strikingly robust fibronectin organization. In this new matrix assembly mechanism, α5β1 integrin-based focal adhesions slide actively on the underlying matrix towards the ventral cell center through the dynamic shortening of myosin IIA-associated actin stress fibers to drive rapid fibronectin fibrillogenesis distal to the adhesion. This mechanism contrasts with classical fibronectin assembly based on stable/fixed-position focal adhesions containing αVβ3 integrins plus α5β1 integrin translocation into proximal fibrillar adhesions. On basement membrane components, these sliding focal adhesions contain standard focal adhesion constituents but completely lack classical αVβ3 integrins. Instead, peripheral α3β1 or α2β1 adhesions mediate initial cell attachment, but over time are switched to α5β1 integrin-based sliding focal adhesions to assemble fibronectin matrix. This basement membrane-triggered mechanism produces rapid fibronectin fibrillogenesis, providing a mechanistic explanation for the well-known widespread accumulation of fibronectin at many organ basement membranes.

## Introduction

Cells secrete and organize multiple types of extracellular matrix to provide structural and biochemical support to cells and tissues (Hay, 1991; McKeown-Longo and Mosher, 1983). Two major structural forms of extracellular matrix are basement membrane and interstitial matrix (Alberts et al., 2015; Jayadev and Sherwood, 2017; Manabe et al., 2008; Pozzi et al., 2017). The matrix glycoprotein fibronectin (FN) is distributed ubiquitously in interstitial matrix. However, FN is also characteristically concentrated immediately adjacent to basement membranes in many adult organs (Aplin and Campbell, 1985; Couchman et al., 1979; Darribere et al., 1986; Gardiner et al., 1986; Gil and Martinez-Hernandez, 1984; Hynes, 1990; Kohno et al., 1987; Leppi et al., 1982; Linder et al., 1978; Mosher, 1989; Uscanga et al., 1984; Warburton et al., 1981) and during embryonic development, where it can function to promote embryonic cell migration (Duband and Thiery, 1982a, b; Krotoski et al., 1986; Mayer et al., 1981; Meier and Drake, 1984; Newgreen and Thiery, 1980). The spatial accumulation of FN at basement membranes was discovered more than four decades ago (Linder et al., 1978; Wartiovaara et al., 1978), but the mechanism remains unclear. We hypothesized that basement membranes might govern FN organization and deposition by direct contact regulation. We tested this hypothesis and discovered a mechanism involving integrin switching and a novel process in which focal adhesions sliding can produce robust local FN fibrillogenesis.

The process of FN matrix assembly is a cell-mediated process following the binding of soluble FN to integrins on the cell surface (Hynes, 1990; Pankov and Yamada, 2002; Singh et al., 2010). Soluble FN dimers exist in a folded, closed conformation (Zaidel-Bar and Geiger, 2010), and integrins are thought to unfold this soluble FN and trigger its polymerization into fibers by applying intracellular forces to FN, transmitted through interactions of their cytoplasmic domains with the actin-myosin cytoskeleton (Hynes, 1999; Schwarzbauer and Sechler, 1999; Zhong et al., 1998). Integrin α5β1 is the major receptor for binding soluble FN and initiating fibril formation (Akiyama et al., 1989; Fogerty et al., 1990; Giancotti and Ruoslahti, 1990). α5β1 binds to the RGD and synergy regions of FN, both of which are essential for de novo FN fibril formation (Nagai et al., 1991; Sechler et al., 1997). Nevertheless, other RGD-binding integrins, such as αVβ3 (Wennerberg et al., 1996; Wu et al., 1996) and αIIbβ3 (Hughes et al., 1996; Wu et al., 1995b), can assemble FN after stimulation, permitting FN matrix formation by α5-null cells (Yang et al., 1993). Some integrins, such as α3β1 (Carter et al., 1990) and α4β1 (Sechler et al., 2000), can bind to non-RGD regions of FN, and α3β1 was reported to promote FN assembly in α5-deficient CHO cells, although the binding of α3β1 to FN is very weak (Wu et al., 1995a).

The main cellular structures that transmit contractile force generated by the actin cytoskeleton to FN are the cell adhesions induced by integrin clustering after binding to FN and other ECM proteins (Singh et al., 2010). These adhesions are contractility-dependent and classified into two major types – focal adhesions and fibrillar adhesions – based on their location, morphology, and molecular composition (Geiger et al., 2001). Classical focal adhesions are oblong cell-attachment structures located mainly at the cell periphery, and they contain primarily αVβ3 and some α5β1 integrins. Focal adhesions also contain a large variety of cytoplasmic molecules that can be classified into several categories, including adaptor proteins (e.g., paxillin, zyxin), actin-associated proteins (e.g., α-actinin, filamin, talin, vinculin), tyrosine kinases (e.g., focal adhesion kinase (FAK) and Src), and phosphatases (e.g. SHP1, PTP-PEST) (Geiger and Yamada, 2011; Horton et al., 2016; Winograd-Katz et al., 2014; Zaidel-Bar and Geiger, 2010). Fibrillar adhesions, in contrast, are very elongated, thread-like structures distributed toward the center of the cell. Fibrillar adhesions are associated closely with FN fibrils (Geiger and Yamada, 2011); they contain integrin α5β1 and tensin, with reduced amounts of focal adhesion components.

Using live-cell imaging and antibody pulse-chase methods, we and others had previously characterized the dynamics of adhesions in vitro, particularly the association of fibrillar adhesion formation with FN fibrillogenesis (Ohashi et al., 2002; Pankov et al., 2000; Zamir et al., 2000). While the vitronectin receptor, αVβ3, remains relatively positionally stable in focal adhesions and firmly anchors cells to the substrate, FN-bound α5β1 integrin molecules continuously translocate from focal adhesions at the cell periphery into fibrillar adhesions. This translocation occurs parallel to actin stress fibers and is thought to transmit tensile forces generated by the actin cytoskeleton to extracellular FN to slowly induce FN fibrillogenesis. The mechanisms of this FN fibrillar assembly model, however, remain somewhat elusive because the exact mechanism of α5β1 integrin translocation is not clear, though it was reported to depend on the actomyosin system (Geiger et al., 2001).

Here, we report that basement membrane and its major components, collagen IV and laminin (Domogatskaya et al., 2012; Fidler et al., 2018), can rapidly induce robust FN assembly, which starts immediately after cell attachment to basement membrane components. Unexpectedly, cells adhering to basement membrane substrates fail to display the classical pattern of FN-assembling adhesions involving stable αVβ3-based focal adhesions and α5β1 integrins translocating along fibrillar adhesions. Instead, intact, actively sliding focal adhesions translocate towards the ventral cell center in association with shortening actin stress fibers by a process requiring myosin IIA in a winch-like mechanism mediating rapid, extensive FN assembly distal to the adhesion. Although these centrally translocating focal adhesions have similar cytoplasmic constituents as regular focal adhesions, they do not contain the αVβ3 integrin, and instead use α2β1 or α3β1 to initiate cell adhesion and fibrillogenesis. As the adhesions mature, α2β1 or α3β1 integrins are then replaced by α5β1 integrins to rapidly assemble FN matrix in a novel integrin-switching process. Taken together, our work describes a novel type of dynamic cell-basement membrane matrix-assembly mechanism that produces rapid, extensive FN fibrillogenesis in a process that can explain the characteristic pattern of FN matrix deposition on basement membranes.

## Results

### Basement membrane and its components strongly induce fibronectin matrix production by cells

FN is strongly enriched at sheet-like basement membranes throughout embryonic development and in many adult organs. It can also promote tumor cell invasion across Matrigel basement membrane-like barriers (Meng et al., 2009). Matrigel is a basement membrane matrix extract from the Engelbreth-Holm-Swarm (EHS) mouse sarcoma rich in basement membrane components such as laminin, collagen IV and heparan sulfate proteoglycan (perlecan) (Kleinman and Martin, 2005). These previous findings suggested the possibility that basement membranes could directly promote FN accumulation. We first verified these scattered prior reports in vivo during mouse development and an in vitro invasion system. FN was readily immunolocalized to the outer surface of basement membranes, e.g., in mouse embryonic lung (Figure S1A). We further confirmed that blocking FN as well as its major cell surface receptor, integrin α5β1, can inhibit in vitro invasion of an oral cancer cell line, SCC-9, through Matrigel (Figures S1B and S1C).

We next tested whether ex vivo basement membranes could directly regulate FN assembly by cultured cells. Intact basement membranes obtained from mouse amniotic membrane strongly and dramatically induced FN fibrillar matrix production by SCC-9 cells (Figures 1A, 1B, S1A and S1D). This phenomenon could be mimicked by seeding cells on growth factor-reduced Matrigel, as well as on each of two purified major basement membrane components: laminin and collagen IV, but not on perlecan nor on the non-basement membrane extracellular matrix proteins vitronectin and collagen I (Figure 1C). Analysis of FN matrix fibers generated on basement membrane, collagen IV, and laminin revealed enhanced FN deposition: fibers were more numerous and were considerably larger in size compared to control cells plated on glass (Figures 1D, 1E and 1F).

**Figure 1.**
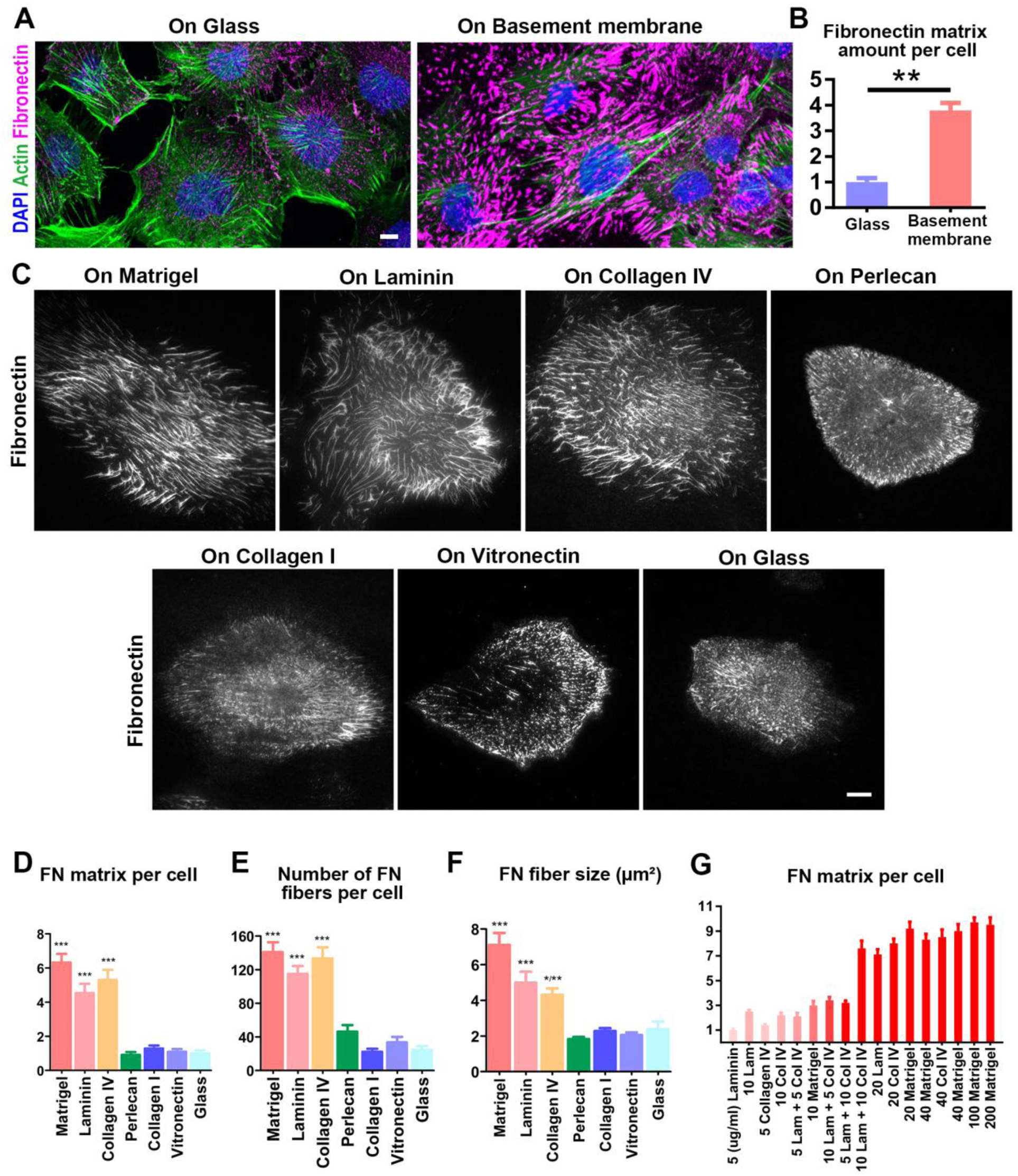
Basement membrane and its major components induce cellular fibronectin matrix deposition. (A) Maximum-intensity projections of immunofluorescence confocal images of SCC-9 cells seeded on glass or basement membrane overnight. (B) Quantification of results in (A) showing FN fibrillar matrix deposition per cell normalized to data from cells on glass. *n*=20. (C) Total internal reflection fluorescence (TIRF) images of SCC-9 cells sparsely seeded on glass or various extracellular matrix protein coatings at 20μg/ml for 8hrs. (D-F) Quantification of results from C showing normalized (to glass) total amounts of FN fibrillar matrix deposition, number of FN matrix fibers per cell, and average area of each FN matrix fiber on different substrates. *n*≥13. p ≤ 0.001 was achieved in all comparisons between each of the conditions of Matrigel, laminin and collagen IV with each of the conditions of collagen I, vitronectin and glass, except in the comparison of FN matrix fiber size between collagen IV and vitronectin (**p ≤ 0.01) and glass (*p ≤ 0.05). (G) Quantification of normalized average FN matrix deposition by each cell growing on various concentrations and combinations of Matrigel, laminin and collagen IV. *n*≥30. Error bars: SEM. *p ≤ 0.05, **p ≤ 0.01, ***p ≤ 0.001. Scale bars: 10 μm. See also Figure S1 and S2.

Importantly, adding soluble Matrigel into culture media instead of coating it onto substrates failed to stimulate FN matrix production, indicating that the basement membrane or its components must be immobilized and be a part of the underying two-dimensional substrate to induce FN assembly (Figure S2A). We tested whether this induction of FN matrix assembly by basement membrane and its components was a general phenomenon shared by other cell types. Various other cancer cell lines, as well as normal human and mouse fibroblasts, showed enhanced fibrillar FN accumulation on Matrigel, although the effect on normal mammary gland epithelial cells (MCF 10A) was marginal (Figure S2B).

Because collagen IV and laminin could each induce robust FN fibrillogenesis, we tested whether these matrix components acted additively or cooperatively by culturing the cells on substrates with varying concentrations and/or combinations of collagen IV and laminin (Figure 1G). Collagen IV and laminin, as well as Matrigel, all achieved maximal FN assembly at approximately 20 µg/ml of protein used for coating substrates. Below this 20 µg/ml protein threshold, the amount of FN matrix assembly was proportional to the concentration of coating. Collagen IV appeared to be a slightly stronger inducer of FN assembly compared to laminin in the SCC-9 cell line. Importantly, when mixed together, collagen IV and laminin had additive but not synergistic effects on FN matrix production.

### Basement membrane promotes efficient fibronectin matrix assembly independent of new protein synthesis or global intracellular signal transduction

To establish whether the induction of FN matrix production by cells is due to increased secretion of soluble FN dimers or to enhanced ability to assemble soluble FN dimers into insoluble FN matrix, we turned to the breast cancer cell line MDA-MB-231, which secretes only very minimal amounts of FN and does not make recognizable FN matrix fibers when cultured in FN-free medium (Figure 2A). When extra FN was added to the medium (20 µg/ml), FN punctate aggregates formed on the cell surface, yet FN fibers were rarely visible. This finding indicates that these cells are not normally capable of assembling a fibrillar FN matrix even when provided with sufficient exogenous FN. Matrigel alone was not able to induce FN matrix production by these cells in FN-free medium, but when extra FN was provided, cells generated substantial quantities of matrix fibers (Figures 2A and 2B). These experiments provided an initial suggestion that Matrigel promotes FN matrix production not by increasing cellular FN secretion, but by enhancing a cell’s capacity to assemble FN matrix – either using FN produced by the cells, or by stimulating the use of exogenous FN.

**Figure 2.**
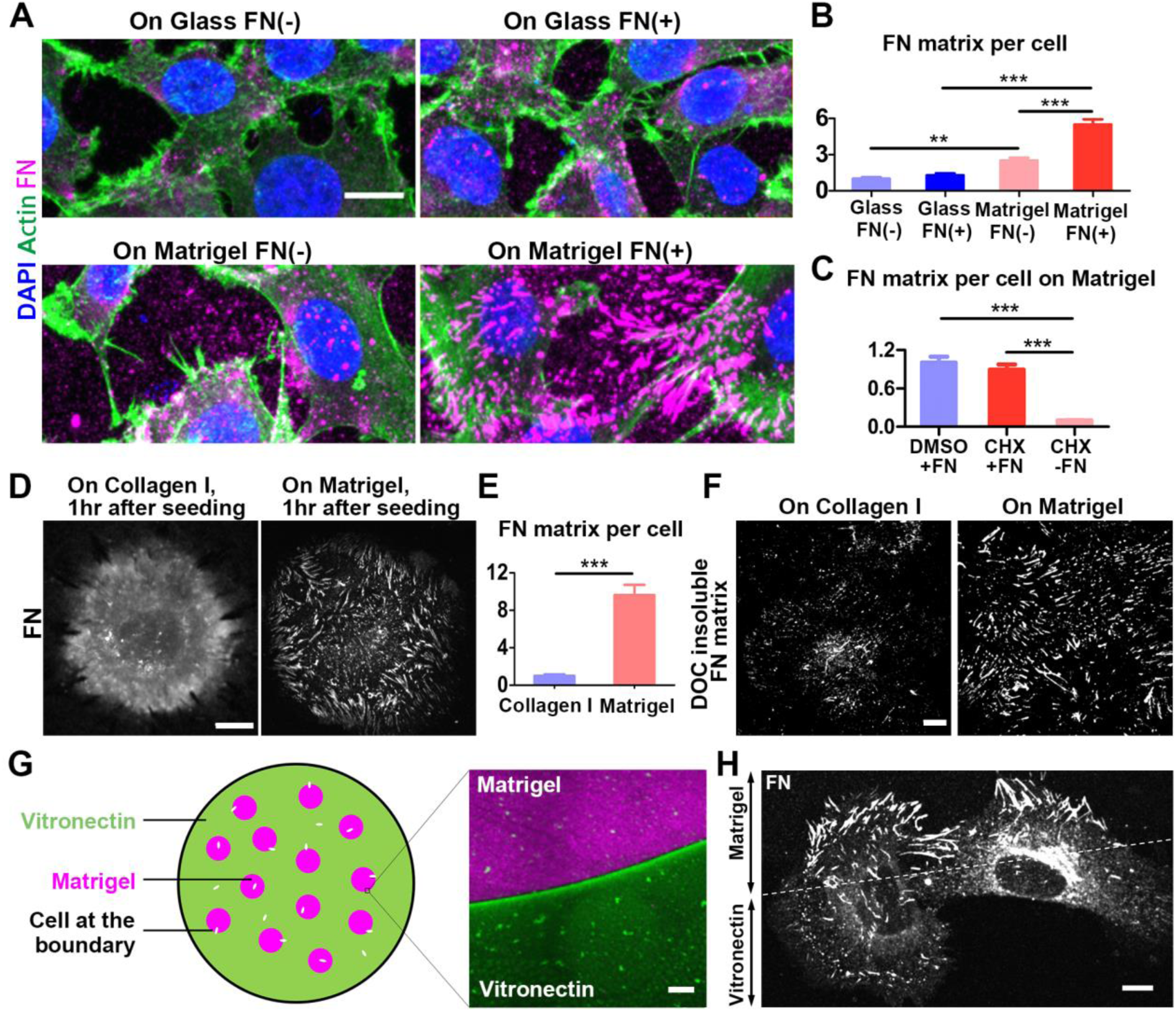
Basement membrane stimulates efficient FN matrix assembly. (A) Maximum-intensity projections of immunofluorescence confocal images of MDA-MB-231 cells seeded on glass or Matrigel with or without exogenous FN (20μg/ml) in the medium. (B) Quantification of results from A showing normalized FN fibrillar matrix deposition per cell. *n*≥15. (C) Quantification of normalized FN matrix deposition per SCC-9 cell cultured in FN-containing medium (20μg/ml) containing DMSO vehicle, FN-containing medium containing cycloheximide (20μg/ml) or FN-free medium containing cycloheximide. *n*=20. (D) Maximum-intensity projections of immunofluorescence confocal images comparing assembly of FN matrix fibers at the bottom of SCC-9 cells seeded on collagen I or Matrigel for 1hr. (E) Quantification of the amount of FN matrix from images shown in D. *n*=12 and 22 for number of cells on collagen I and Matrigel, respectively. (F) Deoxycholate-insoluble FN matrix fibers. Cells growing on Matrigel or collagen I for 1hr were incubated with 10% deoxycholic acid for 10mins before washing and staining with anti-FN. (G) Schematic diagram of the Matrigel-vitronectin patterned coating experiment. (H) Maximum-intensity projections of immunofluorescence confocal images showing FN fibrillogenesis by SCC-9 cells growing at the boundary between Matrigel and vitronectin. Error bars: SEM. **p ≤ 0.01, ***p ≤ 0.001. Scale bars: 10 μm (A, D, F, H); 40 μm (G).

An alternative approach to document a mechanism of FN matrix assembly independent of secretion was to pretreat SCC-9 cells with cycloheximide (20 µg/ml for 4 hrs) to halt FN synthesis and to permit secretion of residual FN to empty the endoplasmic reticulum and Golgi. When exogenous purified FN was provided to these cells, a Matrigel substrate induced strong FN matrix assembly similar to cells that synthesized endogenous FN (Figure 2C). These findings are analogous to the MDA-MB-231 cells that synthesize little FN, and they indicate that Matrigel or its components do not act to promote FN matrix assembly by inducing de novo protein synthesis to stimulate FN assembly, but instead induces a robust assembly mechanism.

To determine how rapidly FN matrix assembly can be induced by Matrigel, we seeded SCC-9 cells on either collagen I or Matrigel for only 1 hr prior to fixation and staining for FN. On collagen I, the well-known phenomenon of direct binding of soluble FN binding to collagen I was observed as a diffuse background along with tiny punctate aggregates and very short fibrils near the cell center. In contrast, on Matrigel, cells assembled elaborate patterns of elongated FN fibrils, even within 1 hr, particularly towards the cell edge (Figures 2D and 2E). These short fibrils retained their structure after 10 min incubation in 10% deoxycholate (Figure 2F), indicating that strong noncovalent FN-fibronectin interactions to form the deoxycholate-resistant fibrils characteristic of assembled fibrillar matrix (McKeown-Longo and Mosher, 1983) had been achieved were present at early stages of FN matrix assembly on Matrigel but not on collagen I.

To test whether a global intracellular signaling pathway or a local subcellular effect was involved in FN assembly induced by Matrigel, we developed a simple patterned extracellular matrix coating method using Matrigel versus vitronectin. After plating, some cells by chance adhered and spread on the substrate boundary between Matrigel and vitronectin (Figure 2G). We found the region of the cell attached to Matrigel substrate assembled much more fibrillar FN matrix than the part of the cell on vitronectin. In fact, the boundary between the two types of ECM coating marked a clear, sharp boundary between robust FN matrix assembly and minimal assembly (Figure 2H). This result demonstrates that Matrigel-induced FN matrix assembly is a localized phenomenon not induced by alterating whole-cell signaling pathways, but instead by a very local response of each region or part of the cell in contact with each specific type of matrix.

### On basement membrane components, fibronectin matrix is assembled by inward sliding of intact focal adhesions

The classical model of FN matrix assembly demonstrates stationary focal adhesions with medial segregation and translocation of the FN-receptor α5β1 integrin into dynamic fibrillar adhesions to generate FN fibrils. To determine if this basement membrane-based mechanism of FN assembly is similar or not to the classical model, we first compared the immunolocalization patterns of the focal adhesion marker paxillin and FN in SCC-9 cells seeded on Matrigel or collagen IV versus cells on vitronectin and collagen I to represent classical matrix assembly conditions (Figure 3A). Consistent with previous findings, cells on collagen I and vitronectin displayed large peripheral focal adhesions containing strong paxillin staining that transitioned to medial fibrillar adhesions with low amounts of paxillin associated with FN fibers. To our surprise, on Matrigel and collagen IV, large numbers of relatively homogenous, strongly paxillin-positive adhesions were found over the entire ventral cell surface. Compared to the typical focal adhesions seen on collagen I or vitronectin, these adhesions were smaller, thinner, and more elongated, with moderate paxillin staining. These adhesions on Matrigel were proximal to, and partially overlapped with, robust FN fibers that were often oriented distal rather than proximal toward the ventral cell center as in classical FN assembly. Interestingly, on vitronectin, a modest number of small, thin FN fibrils could also be found that had a paxillin “cap,” suggesting that some examples of this novel pattern of FN assembly can exist at low levels on classical substrates, but that this pattern is greatly enhanced when cells grow on basement membrane components.

**Figure 3.**
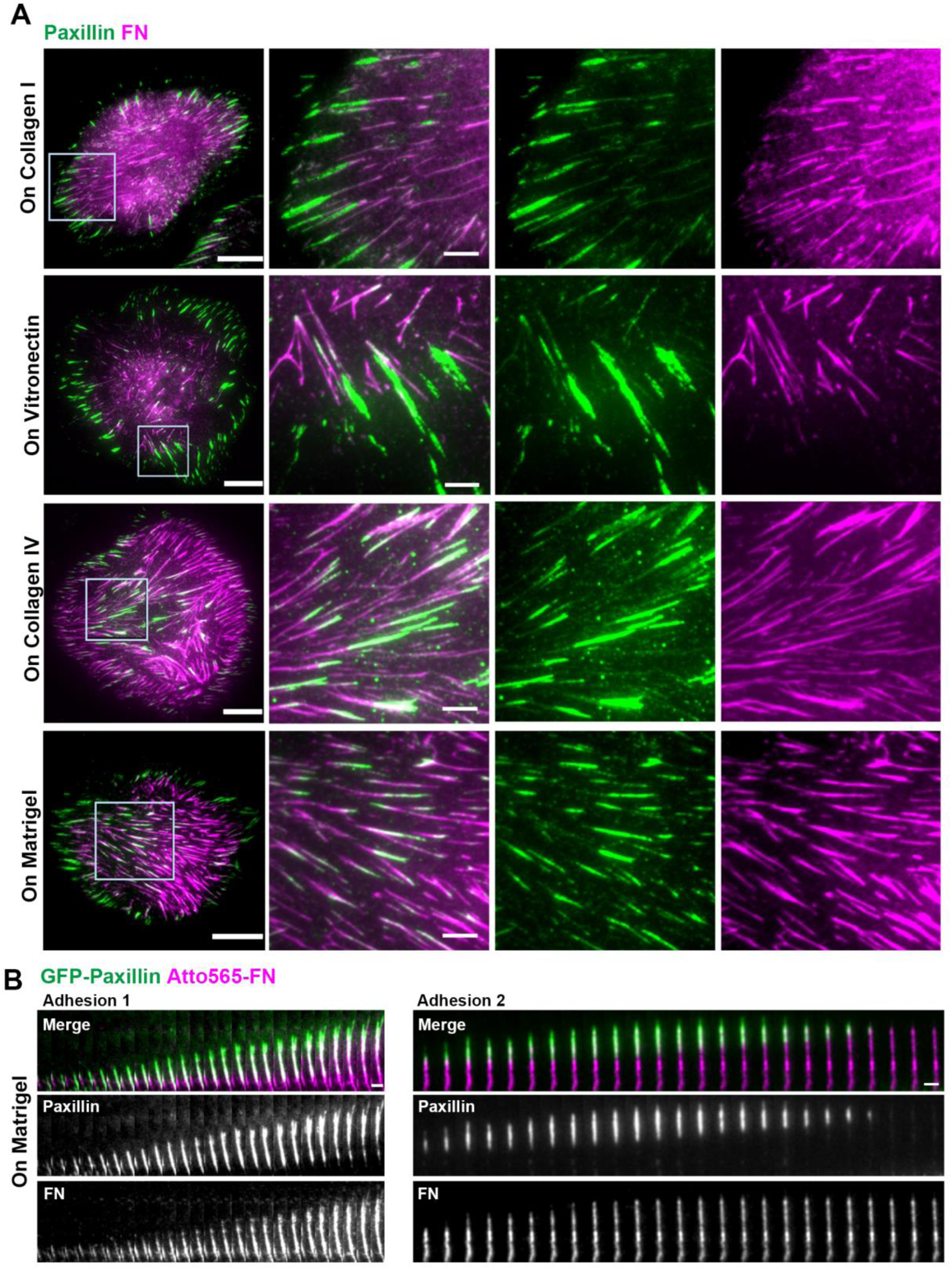
Fibronectin matrix assembly is mediated by sliding focal adhesions on basement membrane. (A) TIRF images of SCC-9 cells seeded on collagen I, vitronectin, collagen IV and Matrigel. Note the patterns of focal adhesions and their spatial relationship with FN fibers highlighted in the insets. (B) Time-lapse image series showing two focal adhesions sliding medially toward the ventral cell center (but shown directed upward for clarity) that are assembling FN matrix fibers. Scale bars: 20 μm (A); 5 μm (insets, B).

To examine the dynamics of these adhesions associated with this alternative FN matrix assembly mechanism, we performed live-cell imaging of GFP-paxillin expressing SCC-9 cells plated on either Matrigel or collagen IV, with soluble Atto565-labeled FN added to the culture medium (Figure 3B and Video S1). Importantly, these focal adhesions, rather than remaining stationery at the cell periphery like classical focal adhesions, translocate or actively slide against the underling matrix as intact focal adhesion structures towards the ventral cell center at speeds similar to classical fibrillar adhesions at 6.4±0.9 μm/hr while generating FN matrix fibers distal to the adhesion. When a focal adhesion was actively sliding, it assembled FN immediately behind it as shown in Figure 3B “Adhesion 1.” The sliding focal adhesion gradually disappeared while the FN fiber associated with it stopped growing and remained stable, as shown in Figure 3B “Adhesion 2.” Interestingly, inward movement of peripheral focal adhesions was reported previously in stationary NIH/3T3 cells growing on glass, but it was not known if it is related to FN fibrillogenesis (Smilenov et al., 1999). Using the same cell line growing on vitronectin, we recapitulated this same pattern of movement and found that while some of the focal adhesions were associated with FN fibers, many were not (Video S2). However, once the cells were placed on Matrigel, focal adhesion movement/sliding and FN fibrillogenesis become tightly coupled to each other (Movie S3). Therefore, Matrigel greatly enhances the association between focal adhesion sliding and FN matrix assembly.

To further characterize this novel class of sliding adhesions, we examined other known focal adhesion markers. These adhesions were positive for not only paxillin, but also for all the focal adhesion components that we examined, including phosphorylated paxillin (Figure 4A), FAK, phosphorylated FAK (Figure 4B), and phospho-tyrosine (Figure 4C). Consequently, these adhesions resemble classical focal adhesions rather than fibrillar adhesions in terms of molecular composition. Interestingly, the signaling-associated molecules phospho-paxillin, FAK, phospho-FAK, and phospho-tyrosine were all strikingly concentrated at the tip of assembling FN fibrils, whereas total paxillin staining was more extended along FN fibrils. This finding suggests that even though these elongated focal adhesions are associated with FN fibers, signaling events may occur only at the inner tip of these translocating focal adhesions as FN is actively assembled. In addition, focal adhesions on Matrigel were also positive for zyxin (Figure 4D), which is regarded as a marker of mature focal adhesions that also serves as a mechanosensor, suggesting that force may be applied to these focal adhesions and possibly contribute to their inward movement and the associated FN fibrillogenesis. Collectively, these data indicate that FN matrix assembly on basement membrane components is mediated by a class of actively inward-sliding focal adhesions.

**Figure 4.**
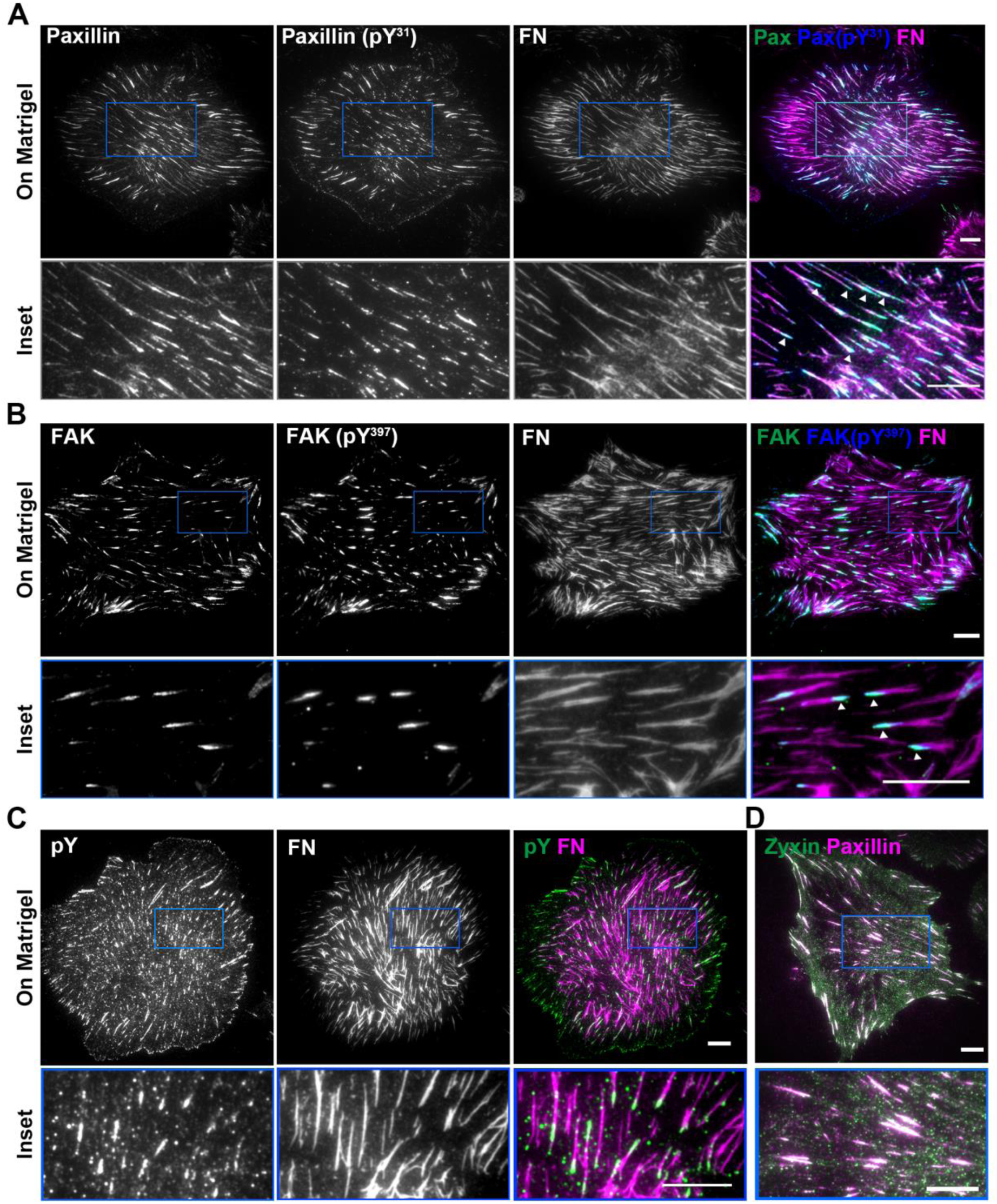
Active fibronectin matrix assembly on basement membrane is associated with medial focal adhesion site rich in tyrosine phosphorylation. TIRF images illustrating the spatial relationship between (A) paxillin, phosphorylated paxillin (Y31) and fibronectin; (B) FAK, phosphorylated FAK (Y397) and fibronectin; (C) phosphotyrosine and fibronectin; (D) zyxin and paxillin; white indicates co-localization. Arrow heads indicate concentration of focal adhesion components on the inner side of FN fibers. Scale bars: 10 μm.

### Inward sliding of focal adhesions on basement membrane components depends on a “contractile winch” system driven by actomyosin contractility

Rho activation and actomyosin contractility are essential for both focal adhesion formation and classical FN fibrillogenesis (Zhong et al., 1998). Actomyosin contractility is a candidate for the force-generating system responsible for the translocation of integrins in fibrillar adhesions. To examine whether Rho activation and actomyosin contractility are also required for this sliding-adhesion FN fibrillogenesis on Matrigel, we treated SCC-9 cells with C3 transferase (Rho inhibitor), Y27632 (Rho-associated protein kinase inhibitor, ROCK inhibitor), or blebbistatin (myosin II ATPase inhibitor) for 3 hours after cell attachment to Matrigel (Figures 5A and S3A). Most of the focal adhesions on Matrigel disappeared after these treatments (Figure S3A). FN fibrillogenesis was substantially impaired by each of these treatments. These results demonstrate that Rho activation and actomyosin contractility are essential for maintaining the sliding focal adhesions on Matrigel and their associated FN fibrillogenesis.

**Figure 5.**
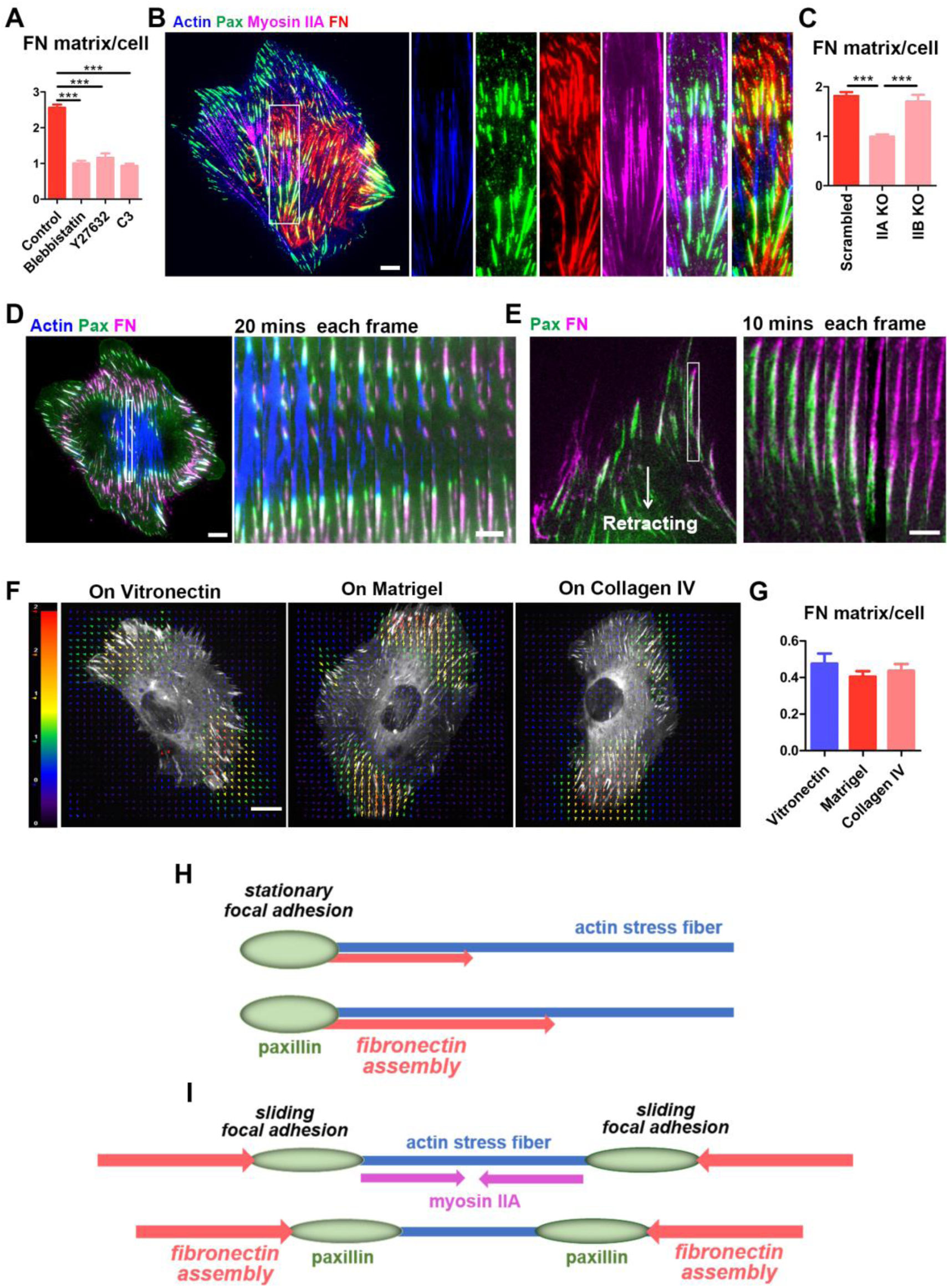
Focal adhesion sliding and fibronectin matrix assembly on basement membrane appears to be driven by an actomyosin winch. (A) Effects of blebbistatin (50μM), Y27632 (20μm), and C3 transferase (4μg/ml) treatment on FN matrix assembly by SCC9 cells growing on Matrigel. *n*≥30. (B) TIRF image of an SCC-9 cell growing on Matrigel. Magnified insets show myosin IIA is associated with actin stress fibers between two sets of parallel focal adhesions. (C) Effects of myosin IIA and myosin IIB knock-out on FN matrix assembly by HFFs on Matrigel. *n*=15. (D) Time-lapse montage of focal adhesions (green) sliding toward each other, showing shortening actin stress fibers (blue) and FN (magenta) matrix fiber elongation. (E) Trailing edge of a cell on Matrigel; white arrow indicates direction of cell movement. Time-lapse montage of the inset shows FN fibrillogenesis (magenta) by retracting focal adhesions (green). (F) Traction force maps of SCC-9 cells growing on vitronectin, Matrigel or collagen IV. (G) Quantification of average traction force magnitude (A.U.). *n*=20. Schematic representation of the classical fibronectin matrix assembly model (H) and the current winch model (I). Error bars: SEM. ***p ≤ 0.001. Scale bars: 10 μm (B, D left, F); 5 μm (D right, E). See also Figure S3.

To evaluate the spatial relationships of contractile actomyosin, focal adhesion sliding, and FN fibrillogenesis on Matrigel, we immunolocalized myosin IIA or IIB isoforms together with FN, actin, and paxillin (Figures 5B and S3B). Overall, myosin IIA was more prominently localized to stress fibers associated with FN fibrils. Intriguingly, the more peripheral adhesions displayed FN fibrils oriented distally towards the periphery, rather than towards the cell interior as during classical FN matrix assembly. Focal adhesions were located at varying distances interior to the cell edge, suggesting that they could have been pulled inward while depositing FN at their rear. Particularly noteworthy were paired focal adhesions, often found in a region of the ventral cell surface between the cell edge and nucleus, facing each other and linked by actin and associated myosin IIA (with little myosin IIB) showing FN deposited at the distal end of each focal adhesion (Figure 5B). Unlike the previous classical FN assembly mechanism postulating motility of FN-receptor integrins and FN along stable actin stress fibers, the spatial patterns we observe – combined with live-cell imaging that documents shrinking of actin stress fibers linking converging focal adhesions (Figure 5D, Videos S4 and S5) – identify a novel phenomenon involving shortening of stress fibers as focal adhesions slide toward each other. In fact, each of two focal adhesions facing each other from the ends of shortening actin microfilament bundles demonstrate a strikingly coordinated translocation toward each other together with shortening of the stress fibers (Videos S4 and S5). These findings point to a model involving a mechanical actomyosin “contractile winch,” where shortening stress fibers linked to focal adhesions mediate the focal adhesion sliding, accompanied by FN assembly (Figure 5I). This model contrasts with the classical FN assembly model in which focal adhesions and stress fibers remain relative stable (Figure 5H).

To distinguish the role of different myosin subtypes in focal adhesion formation and FN fibrillogenesis, we generated myosin IIA and myosin IIB knock-out human fibroblasts. Myosin IIA knock-out, but not myosin IIB knock-out, inhibited focal adhesion formation and FN fibrillogenesis, an effect that was very similar to the effects on SCC-9 cells of treatment by Rho, ROCK, or myosin II inhibitors (Figures 5C and S3C). Therefore, myosin IIA serves as a major force generator for this sliding focal adhesion mechanism of FN fibrillogenesis on basement membrane.

### Distinct focal adhesion pattern on basement membrane components is not associated with altered physical interaction between cell and substrate

Mechanical force is essential for the development and stabilization of focal adhesions, and focal adhesions also serve as major sites for transmission of bidirectional mechanical force between the extracellular matrix and an interacting cell. Consequently, an alternative to the mechanical model of a contractile winch system that pulls together focal adhesions that face each other at ends of actomyosin bundles might be some type of active focal adhesion traction/translocation system in which focal adhesions would exert traction and locomotory forces on the substrate to translocate actively towards the cell center. We used traction-force microscopy to search for such traction forces potentially used by translocating focal adhesions. Instead, we found that cells have similar levels and patterns of contractility on Matrigel, collagen IV, and vitronectin (Figures 5F and 5G). Moreover, even though focal adhesions slide along the ventral cell surface on Matrigel and collagen IV, while remaining stationary at the cell edge on vitronectin, cellular contractile forces were all distributed along the cell edge where classical focal adhesions exist, regardless of the type of substrate (Figure 5F) – we could find no evidence for higher traction forces near the cell center associated with the regions where sliding focal adhesions actively translocate on Matrigel and collagen IV. These data indicate that a basement membrane substrate does not alter focal adhesion traction force patterns or dynamics, consistent with the alternative model of stress-fiber actomyosin “contractile-winch” linking sliding focal adhesions.

Cells often retract entire cell edges during cell migration and when changing direction. Such retractions of large regions of the cell likely generate shear forces on focal adhesions and the substrate. Interestingly, we found as cells on Matrigel retract their edges, focal adhesions at that cell edge slide inward and generate very striking patterns of FN matrix assembly behind them as they slide (Figure 5E and Video S6). This enhanced form of FN matrix assembly on Matrigel involving the sliding of multiple retracting focal adhesions is conceptually complementary to, but often more dramatic than, the local sliding of single focal adhesions in the winch-like mechanism described earlier. This retraction-associated sliding of focal adhesions that produces large amounts of FN fibrils can also provide an explanation for the frequent presence of FN fibrillogenesis adjacent to cell edges on substrates consisting of basement membrane components (e.g., see Figure 1C).

### Integrin α5β1 is required for focal adhesion sliding but is only partially essential for fibronectin matrix assembly on Matrigel

Integrin α5β1 is the primary integrin responsible for FN matrix assembly under classical conditions and is the only integrin pair found in fibrillar adhesions. To investigate the role of α5β1 in FN fibrillar matrix assembly on Matrigel, we first immunolocalized α5β1 compared to paxillin and FN in cells grown on Matrigel for 6 hrs (Figure 6A). Both α5β1 and paxillin were co-distributed along FN fibers; along a single fiber, α5β1 was often associated with the inner (medial) end of the fibrillar bundle, and in some cases both α5β1 and FN were located slightly behind paxillin. Therefore, α5β1 integrin is present in the sliding focal adhesions on Matrigel, and is a candidate as a key component linking contractile force from the contractile-winch model for FN fibrillogenesis.

**Figure 6.**
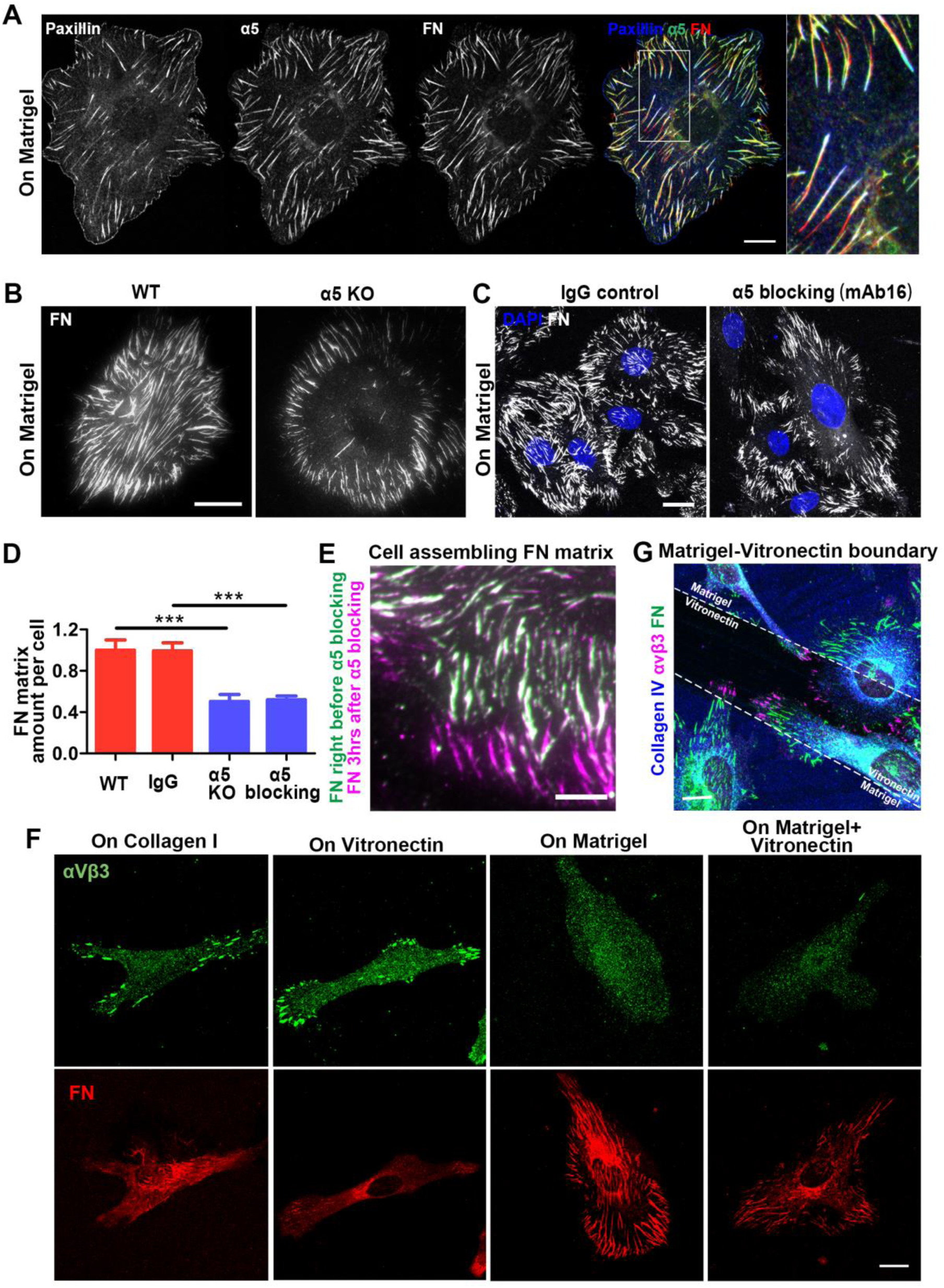
Winch-mediated fibronectin matrix assembly is partially dependent on integrin α5β1, and does not require integrin αVβ3. (A) Confocal images of an SCC-9 cell seeded on Matrigel for 6 hrs and immunostained for paxillin, integrin α5β1, and fibronectin. Remaining fibronectin matrix fibers form primarily near the cell periphery after integrin α5β1 blocking (B) or knock-out (C). (D) Quantification of fibronectin matrix of results from B and C. *n*=15 for blocking, *n*=20 for knock-out. (E) Merged TIRF live imaging of assembled fibronectin matrix fibers immediately before integrin α5β1 blocking and 3hrs after blocking; note the new fibronectin matrix fibers (magenta) assembled on the cell edge after blocking. (F) Integrin αVβ3 aggregation in focal adhesions of cells on vitronectin or collagen I was lost on Matrigel and could not be rescued by mixing Matrigel with vitronectin for coating the substrate. (G) Cells growing at the boundary between Matrigel and vitronectin. Note that within a single cell, only the region of the cell body on vitronectin shows integrin αVβ3 clustering, while the regions on Matrigel are more effective at assembling fibronectin matrix than on vitronectin. Error bars: SEM. ***p ≤ 0.001. Scale bars: 10 μm (A, E, F); 20 μm (B, C, G). See also Figure S4.

β1 integrin activity can be enhanced by MnCl_2_ or activating anti-β1 (clone TS2/16) and facilitates classical FN matrix assembly. However, the monoclonal 9EG7 antibody specifically targeting activated β1 revealed similar level of brightness in staining the fibrillar adhesions on glass and sliding focal adhesions on Matrigel (Figure S4), suggesting the contractile winch is not promoted by increased β1 activity.

To determine the role of α5β1 integrin in Matrigel-induced FN fibrillogenesis, we ablated the α5β1 integrin using CRISPR-Cas9. Cells lacking this FN receptor lacked the pattern of substantially enhanced FN assembly towards the cell center distinctively associated with the sliding focal adhesions of the contractile winch. Nevertheless, cells could still generate some FN assembly at the cell periphery (Figure 6B). A similar effect was achieved using blocking antibodies against α5β1 integrin (Figures 6C and 6D). In fact, during live imaging, FN matrix fibers stopped extending after α5β1 integrin blocking, but the cell could still generate new FN fibers at the cell edge (Figure 6E). This observation indicates that α5β1 integrin is responsible for FN fibrillogenesis during focal adhesion sliding; however, non-α5β1 integrin(s) also participate and partially compensate for α5β1 loss in FN matrix assembly on Matrigel, particularly near the cell periphery.

In the classical FN matrix assembly model, αVβ3 is the integrin that remains in focal adhesions, providing stable anchorage to cells as α5β1 translocates inward for FN fibrillogenesis. Previous knockout studies of α5β1 have established that αVβ3 can also bind the RGD region of FN and mediate its assembly (Wu et al., 1996). Therefore, we tested whether the αVβ3 integrin could serve as the non-α5β1 integrin and contribute to FN matrix assembly by cells on Matrigel. Immunostaining of αVβ3 in SCC-9 cells grown on different substrates (Figure 6F) disproved this hypothesis. On vitronectin and collagen I, αVβ3 was present in the focal adhesions on the cell edge as previously described. On Matrigel, to our surprise, αVβ3 was only diffusely distributed throughout the plasma membrane, with no concentration in focal adhesions. A theoretically possible explanation for this phenomenon could have been that Matrigel coating prevents vitronectin binding to glass so that it cannot be used by cells to attach and induce αVβ3 aggregation in focal adhesions. Therefore, we cultured cells on glass coated with a mixture of vitronectin and Matrigel. However, once again, αVβ3 failed to concentrate in focal adhesions even when vitronectin was available on the substrate, suggesting that αVβ3 is inhibited by Matrigel. In fact, similar to its local promotion of FN matrix assembly (Figure 2H), Matrigel fails to promote αVβ3 in focal adhesions in the region of the cell contacting Matrigel, even though other parts of the same cell attaching to vitronectin readily generate classical focal adhesions (Figure 6G). This finding is again consistent with a local response to specific basement membrane components, rather than whole-cell signaling.

### Focal adhesion sliding is enabled by a α3β1-to-α5β1 integrin switch

We next determined the non-α5β1 integrin that could mediate FN matrix assembly on Matrigel. Immunofluorescence screening revealed that the α3β1 integrin co-localizes with the peripheral FN matrix fibers after α5β1 blockade (Figure 7A). α3β1 integrin is a well-known laminin receptor, and its interaction with collagen IV has also been reported (Miles et al., 1995; Takada et al., 1991). In addition, α3β1 can assemble FN matrix in α5-deficient CHO cells (Wu et al., 1995a). Interestingly, in cells fixed 3-hours after plating, α3β1 did not aggregate in focal adhesions when α5 integrin was not blocked (Figure 7A), indicating that it acted in a compensatory manner to the absence of the α5 integrin. Moreover, cycloheximide-treated SCC-9 cells used α3β1 to attach to Matrigel in FN-free medium (data not shown). Therefore, we hypothesized that FN matrix assembly on Matrigel might be a two-step process in which α3β1 mediates initial cell attachment and early FN matrix assembly, and then α5β1 localizes later to suppress and supplant α3β1 while mediating the large induction of FN matrix assembly.

**Figure 7.**
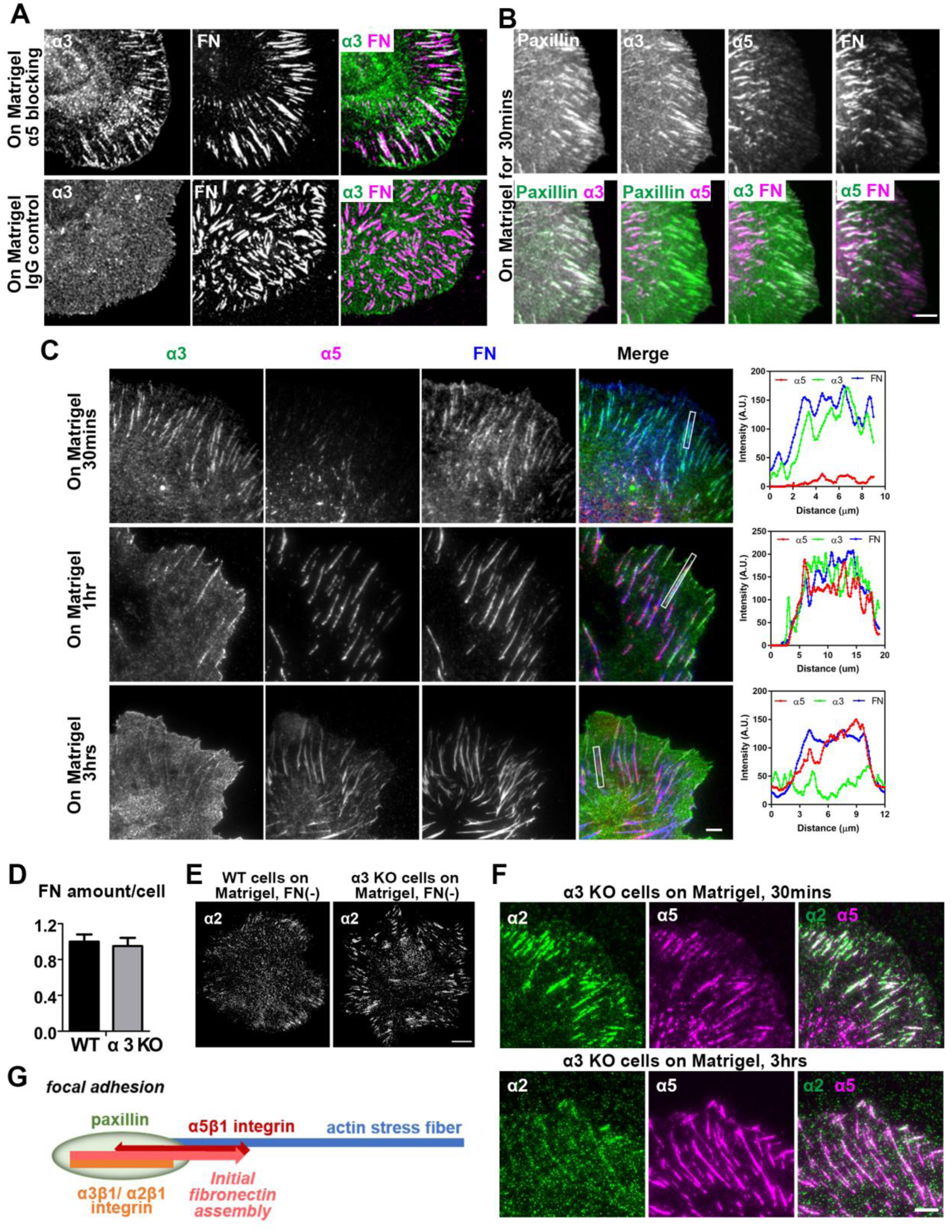
The contractile winch is triggered by an integrin switch from α3β1/α2β1 to α5β1. (A) Diffuse distribution of integrin α3β1 on Matrigel and its aggregation in adhesions and association with fibronectin matrix fibers after α5β1 blocking. (B) TIRF images of SCC-9 cells seeded on Matrigel for 30mins and immunostained for paxillin, integrin α3β1, integrin α5β1, and fibronectin. (C) TIRF images and plot profiles of SCC-9 cells seeded on Matrigel for 30mins, 1hr, and 3hrs and immunostained for integrin α3β1, integrin α5β1, and fibronectin. Note the gradual disappearance of α3β1 and the appearance of α5β1 immunolocalization. (D) Quantification reveals no difference in fibronectin matrix assembly by wild-type cells and α3β1 knock-out cells. *n*=30. (E) Increased aggregation of integrin α2β1 in cell-Matrigel adhesions after α3β1 knock-out. (F) TIRF images illustrating α5β1 integrin becoming the dominant integrin against α2β1 integrin from 30mins to 3hrs after plating. (G) Schematic drawing of integrin switching in the focal adhesion that eventually triggers winch-mediated FN matrix assembly. Error bars: SEM. Scale bars: 10 μm (A, E); 5μm (B, C, F).

To elucidate the temporal dynamics of α3β1 involvement in initial FN matrix assembly by SCC-9 cells on Matrigel, we plated these cells on Matrigel for only 30 mins and then immunostained for paxillin, α3β1, α5β1, and FN (Figure 7B). Intriguingly, at this early time point, α3β1 was found at the cell edge in early focal adhesions colocalized with paxillin, while α5β1 was located on the inner (medial) side of α3β1. Both integrins were associated with FN, although the FN fibers associated with α5β1 were more prominent (brighter by immunostaining) than those associated with α3β1. Therefore, the initial step of FN matrix assembly on Matrigel appears to be similar to the classical model of assembly: α3β1 is localized in stable peripheral focal adhesions (similar to αVβ3 in the classical model), while α5β1 localizes to the inner side of focal adhesions in a position to mediate FN fibrillogenesis, analogous to its role in fibrillar adhesions in the classical model.

To determine how this initial mode of FN matrix assembly evolves into the sliding focal adhesion mode of enhanced assembly, we characterized the dynamics of both integrins compared to FN in cells plated on Matrigel for 30 mins, 1 hr and 3 hrs, then fixed and immunostained (Figure 7C). The α3β1 integrins in the peripheral focal adhesions were gradually replaced by α5β1: at approximately 1 hr after cell seeding, α5β1 was present in focal adhesions together with α3β1. By 2 hours later, α5β1 occupied the focal adhesions, while α3β1 was barely detectable. This switch of integrins may to be due to their affinities for FN: α5β1 might have a higher affinity for the FN initially assembled by α3β1 and gradually replace α3β1 in focal adhesions. We suggest that once α5β1 becomes the dominant integrin in focal adhesions, the winch system can begin operating to assemble FN matrix fibers efficiently.

### α2β1 integrin can compensate and substitute for the α3β1 integrin to initiate the integrin switch

To test the role of α3β1 in FN matrix assembly on Matrigel, we generated α3β1 CRISPR-Cas9 knock-out SCC-9 cells. These cells could still assemble similar amounts of FN matrix on Matrigel compared to wild-type cells (Figure 7D). Consequently, the proposed “winch” mechanism does not depend solely on preceding α3β1 function. Since cycloheximide-treated α3β1 knock-out cells can still attach to Matrigel and form focal adhesions in FN-free medium (data not shown), we speculated that another integrin was substituting for the function of α3β1. Therefore, we performed another round of integrin screening and found that α2β1 was upregulated and enriched in focal adhesions in α3β1 knock-out cells initially attached to Matrigel (Figure 7E). We further demonstrated that similar to α3β1, the α2β1 integrin was later switched to α5β1 (Figure 7F). Therefore, the α2β1 integrin can substitute for α3β1 for initial focal adhesion assembly and can precede the introduction of α5β1 into focal adhesions, promote initial FN assembly, and participate in the integrin switch to α5β1 (Figure 7G).

## Discussion

The major finding of this study is that basement membrane – and its major components as substrates – can dramatically induce enhanced fibronectin matrix assembly in a variety of cell types. This novel form of matrix assembly is mediated by distinct focal adhesions to basement membrane components that slide toward the cell center in a process driven by an actomyosin contractile winch. This winch is activated upon a switch from using receptors for basement membrane components, α2β1 and α3β1, to the FN receptor α5β1. This novel process can explain the long-standing, unexplained observation of FN matrix localization immediately adjacent to basement membranes in vivo, which we find to be driven by a novel integrin-switching, dynamic cell-matrix adhesion mechanism.

The contractile-winch model we propose differs from the classical FN matrix assembly model and the inward movement/sliding of focal adhesions in previous studies (Pankov et al., 2000; Smilenov et al., 1999; Zamir et al., 2000). In the classical model, cell-matrix adhesions segregate into stable peripheral focal adhesions and translocating fibrillar adhesions. Translocation of α5β1 integrin in fibrillar adhesions away from focal adhesions depends on actomyosin contractility, but how contractility and translocation are coupled is not clear (Geiger and Yamada, 2011; Pankov et al., 2000). Similar to focal adhesions, actin stress fibers remain relatively stable during α5β1 integrin translocation (Ohashi et al., 2002). In the contractile-winch model, however, we observe no segregation of cell-matrix adhesions, and instead focal adhesions actively slide inward or toward each other, driven by myosin IIA-mediated shortening of actin stress fibers. The previously described movement of focal adhesions in stationary but not migratory cells might be a related process, but our time-lapse imaging showed that even though sliding can occur on vitronectin, FN assembly requires a substrate of a basement membrane component for contractile-winch α5β1-mediated FN fibrillogenesis. Therefore, this model represents a direct connection between a specific class of protein substrate, actomyosin contractility, focal adhesion sliding, and enhanced FN matrix assembly.

Cell matrix dynamics and FN matrix assembly can be affected by the composition and physical attributes of the substrate. Covalently linking FN to the substrate prevents segregation of cell-matrix adhesions, leading to α5β1 integrin enrichment in paxillin-rich focal adhesions (Katz et al., 2000), resembling the effects of basement membrane components reported here. Rigid substrates facilitate FN matrix assembly (Halliday and Tomasek, 1995). The elasticity of substrates can affect Rho activity and regulate cell contractility to expose cryptic domains for FN-FN binding (Yoneda et al., 2007; Zhang et al., 1999; Zhong et al., 1998). Importantly, even though Rho activity and actomyosin contractility are required for the contractile-winch matrix assembly model, cells do not generate any additional detectable traction force overall, nor at sites where focal adhesions are translocating on basement membrane components. There are to our knowledge no known FN binding sites on laminin and collagen IV, and neither soluble laminin nor collagen IV could bind to assembled FN matrix fibers, nor to urea-unfolded FN (data not shown). Therefore, basement membrane components may not have any special mechanical interactions with FN for activating the contractile winch.

α5β1 is the primary integrin component of both fibrillar adhesions in the classical FN assembly model and sliding focal adhesions in the winch model. In the classical model, αVβ3 integrin binds to vitronectin and remains in focal adhesions to provide anchoring sites necessary for α5β1 integrin translocation. In the contractile-winch model, αVβ3 integrins are notably absent from the focal adhesions on basement membrane components. This absence of αVβ3 is not due to shortage of ligand, since adding vitronectin to substrates also failed to produce αVβ3 accumulation in the focal adhesions. Nevertheless, αVβ3 clustered in focal adhesions at portions of the cell contacting vitronectin, but not Matrigel. It was also not due to global trans-dominant inhibition of integrins, since neither antibody inhibition nor knock-out of α5β1 or α3β1 could restore αVβ3 to focal adhesions (data now shown) (Gonzalez et al., 2010; Gonzalez et al., 2002; Ly et al., 2003).

Instead of αVβ3 in focal adhesions on basement membrane substrates, we identified α3β1, a receptor for both laminin and collagen IV (although the latter is controversial) (Miles et al., 1995; Takada et al., 1991), as mediating both cell attachment to Matrigel and initial focal adhesion formation. Interestingly, α3β1 has been reported to assemble FN matrix in α5-null CHO cells (Wu et al., 1996), consistent with our finding that SCC-9 cells treated with α5-blocking antibodies can still generate some FN fibers that co-localize with α3β1. Importantly, we find that α5β1 integrins are located on the inner (medial) side, but not within, newly formed focal adhesions on Matrigel. Nevertheless, α5β1 integrins manage to move into the pre-existing α3β1-positive focal adhesions and then trigger the contractile-winch mechanism by gradually displacing α3β1, probably through higher affinity to the FN that had been initially assembled by α3β1.

If α3β1 is genetically ablated, we found that α2β1, a well-known collagen IV receptor and laminin receptor (Kern et al., 1993; Languino et al., 1989; Staatz et al., 1990), becomes concentrated in focal adhesions in α3β1 knock-out cells. The α2β1 integrin could in fact substitute for α3β1 to mediate cell attachment, early FN assembly, and the integrin switch to α5β1. Interestingly, α2β1 is also upregulated in α5β1 knock-out cells (but not in cells treated with α5β1 blocking antibodies) and co-localizes with FN (data not shown). In short, the winch-driven focal adhesion sliding mechanism appears to require sequential functions of different integrins to bind to substrate proteins and mediate subsequent local integrin interactions for inducing FN matrix assembly.

We found no role for enhanced β1 activity in the large basement membrane induction of matrix assembly. Intracellular signaling might contribute to the winch-mediated FN matrix assembly mechanism. FAK plays central roles in fibrillar adhesion organization and FN matrix assembly (Ilic et al., 2004). Src kinase contributes to this process, as well as a recently recognized AMPK pathway (Georgiadou et al., 2017; Wierzbicka-Patynowski and Schwarzbauer, 2002). Src kinase can maintain the assembled FN matrix through its phosphorylation of paxillin (Wierzbicka-Patynowski et al., 2007). However, non-phosphorylated paxillin is essential for fibrillar adhesion formation, while phosphorylated paxillin preferentially interacts with FAK and increases focal adhesion dynamics (Zaidel-Bar et al., 2007). In the current contractile-winch model, phosphorylated paxillin and the more general category of phosphotyrosine-containing proteins are notably enriched and concentrated at the inner tip of sliding focal adhesions where FN is being assembled. These data suggest that phosphorylation of paxillin may serve as a downstream effector of various upstream regulators of FN matrix assembly in different assembly mechanisms and at different assembly stages.

Our findings provide a simple explanation for the long-standing observation of FN matrix accumulation at basement membranes in a wide variety of tissues in vivo. During embryonic development, FN deposition along basement membranes provides guidance for cell migration, as in the case of neural crest cells migrating along the basement membrane of neural tube, somites, and ectoderm (Duband and Thiery, 1982b; Krotoski et al., 1986; Mayer et al., 1981; Meier and Drake, 1984; Newgreen and Thiery, 1980). Similarly, mesenchymal cells migrate along FN on basement membranes during avian and amphibian gastrulation (Darribere et al., 1986; Duband and Thiery, 1982a). During coagulation, FN exposure following damage of blood vessels can mediate platelet adhesion to basement membranes (Houdijk et al., 1986). Intriguingly, at least in oral and lung cancer cells, FN can reportedly contribute to cancer cell invasion through basement membrane-like Matrigel barriers (Houdijk and Sixma, 1985; Meng et al., 2009). These observations, combined with our data, suggest that cancer cells might sometimes hijack basement membrane to assemble FN matrix fibrils to facilitate their invasion through basement membranes.

## Acknowledgments

This research was supported by the Intramural Research Program of the NIH, NIDCR, and by fellowship support to Jiaoyang Lu from the China Scholarship Council (CSC) and The National Natural Science Foundation of China (NSFC). We thank Matthew Kutys for the lentiviral CRSPR-Cas9 lentiviruses and the NIDCR imaging core for assistance in image acquisition.

## Author contributions

Conceptualization, J.L. and K.M.Y.; Methodology, J.L, K.M.Y., A.D.D., S.W.; Investigation, J.L, Y.S., M.A.B.; Writing-Original Draft, J.L.; Writing-Review & Editing, K.M.Y., A.D.D., S.W.; Funding Acquisition, K.M.Y. and M.Z.

## METHODS

### KEY RESOURCES TABLE

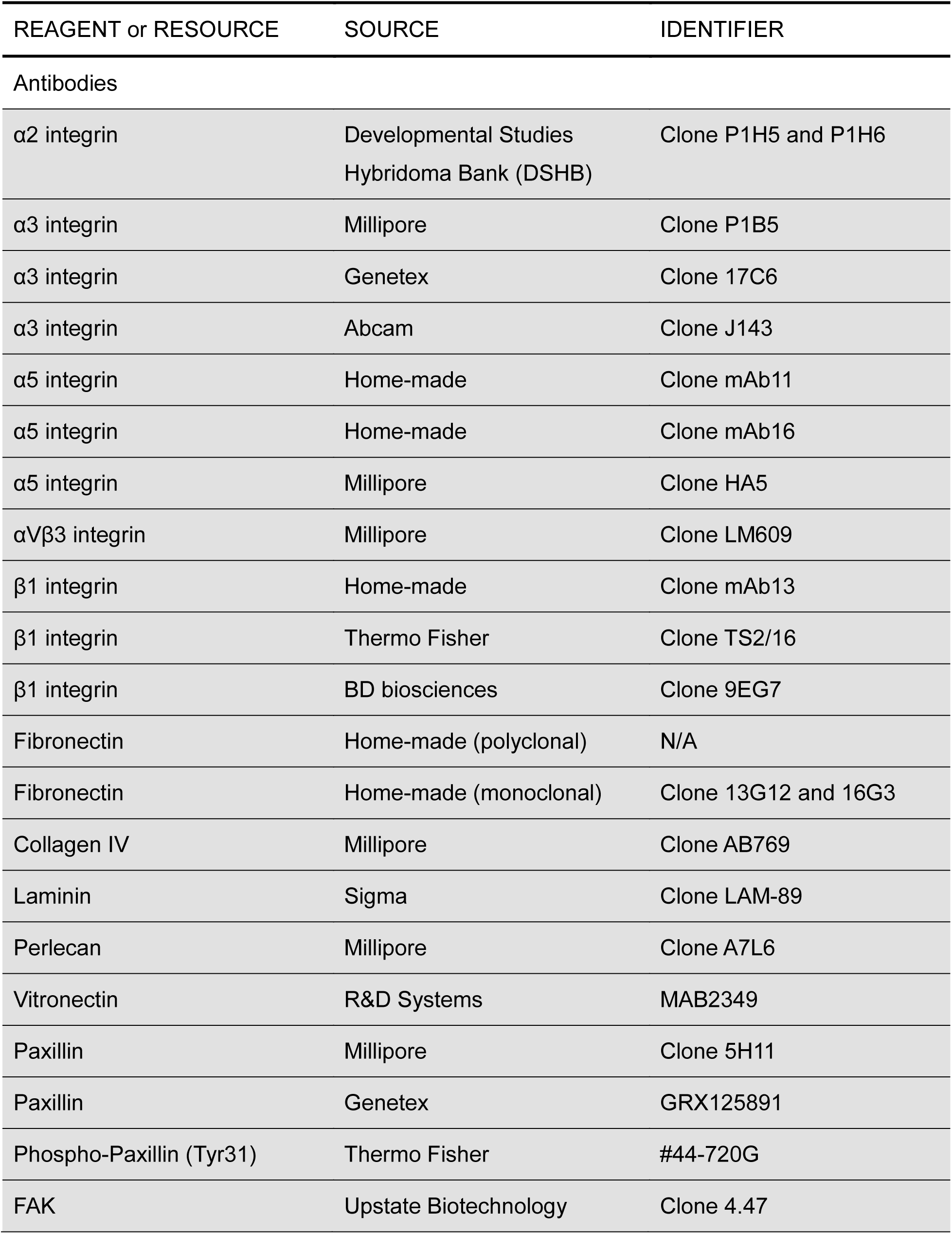
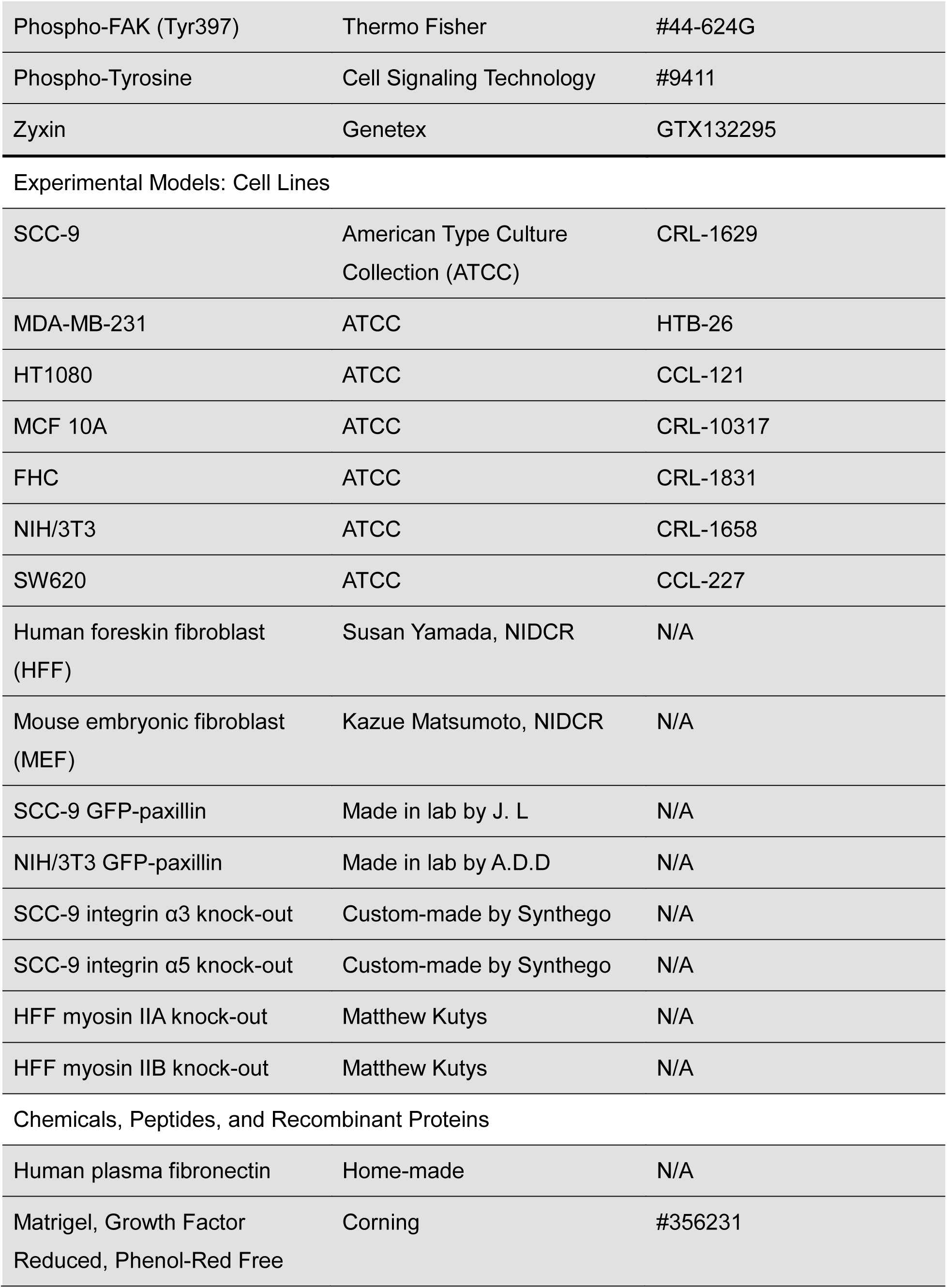
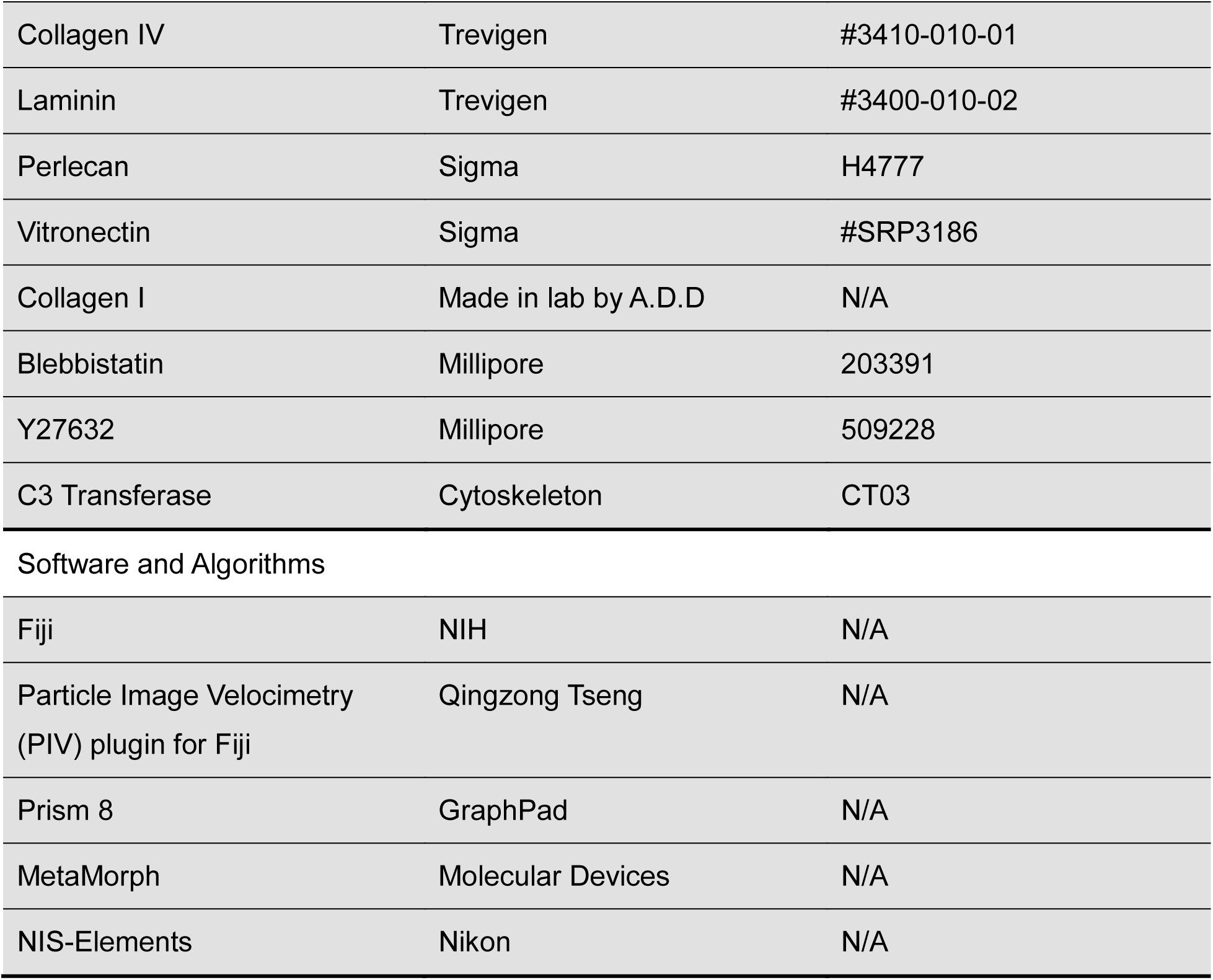

| REAGENT or RESOURCE | SOURCE | IDENTIFIER |
| --- | --- | --- |
| Antibodies |  |  |
| $\alpha 2$ integrin | Developmental Studies Hybridoma Bank (DSHB) | Clone P1H5 and P1H6 |
| $\alpha 3$ integrin | Millipore | Clone P1B5 |
| $\alpha 3$ integrin | Genetex | Clone 17C6 |
| $\alpha 3$ integrin | Abcam | Clone J143 |
| $\alpha 5$ integrin | Home-made | Clone mAb11 |
| $\alpha 5$ integrin | Home-made | Clone mAb16 |
| $\alpha 5$ integrin | Millipore | Clone HA5 |
| $\alpha V\beta 3$ integrin | Millipore | Clone LM609 |
| $\beta 1$ integrin | Home-made | Clone mAb13 |
| $\beta 1$ integrin | Thermo Fisher | Clone TS2/16 |
| $\beta 1$ integrin | BD biosciences | Clone 9EG7 |
| Fibronectin | Home-made (polyclonal) | N/A |
| Fibronectin | Home-made (monoclonal) | Clone 13G12 and 16G3 |
| Collagen IV | Millipore | Clone AB769 |
| Laminin | Sigma | Clone LAM-89 |
| Perlecan | Millipore | Clone A7L6 |
| Vitronectin | R&D Systems | MAB2349 |
| Paxillin | Millipore | Clone 5H11 |
| Paxillin | Genetex | GRX125891 |
| Phospho-Paxillin (Tyr31) | Thermo Fisher | #44-720G |
| FAK | Upstate Biotechnology | Clone 4.47 |
| Phospho-FAK (Tyr397) | Thermo Fisher | #44-624G |
| Phospho-Tyrosine | Cell Signaling Technology | #9411 |
| Zyxin | Genetex | GTX132295 |
| Experimental Models: Cell Lines |  |  |
| SCC-9 | American Type Culture Collection (ATCC) | CRL-1629 |
| MDA-MB-231 | ATCC | HTB-26 |
| HT1080 | ATCC | CCL-121 |
| MCF 10A | ATCC | CRL-10317 |
| FHC | ATCC | CRL-1831 |
| NIH/3T3 | ATCC | CRL-1658 |
| SW620 | ATCC | CCL-227 |
| Human foreskin fibroblast (HFF) | Susan Yamada, NIDCR | N/A |
| Mouse embryonic fibroblast (MEF) | Kazue Matsumoto, NIDCR | N/A |
| SCC-9 GFP-paxillin | Made in lab by J. L | N/A |
| NIH/3T3 GFP-paxillin | Made in lab by A.D.D | N/A |
| SCC-9 integrin $\alpha 3$ knock-out | Custom-made by Synthego | N/A |
| SCC-9 integrin $\alpha 5$ knock-out | Custom-made by Synthego | N/A |
| HFF myosin IIA knock-out | Matthew Kutys | N/A |
| HFF myosin IIB knock-out | Matthew Kutys | N/A |
| Chemicals, Peptides, and Recombinant Proteins |  |  |
| Human plasma fibronectin | Home-made | N/A |
| Matrigel, Growth Factor Reduced, Phenol-Red Free | Corning | #356231 |
| Collagen IV | Trevigen | #3410-010-01 |
| Laminin | Trevigen | #3400-010-02 |
| Perlecan | Sigma | H4777 |
| Vitronectin | Sigma | #SRP3186 |
| Collagen I | Made in lab by A.D.D | N/A |
| Blebbistatin | Millipore | 203391 |
| Y27632 | Millipore | 509228 |
| C3 Transferase | Cytoskeleton | CT03 |
| Software and Algorithms |  |  |
| Fiji | NIH | N/A |
| Particle Image Velocimetry (PIV) plugin for Fiji | Qingzong Tseng | N/A |
| Prism 8 | GraphPad | N/A |
| MetaMorph | Molecular Devices | N/A |
| NIS-Elements | Nikon | N/A |

### CONTACT FOR REGENT AND RESOURCE SHARING

Further information and requests for resources and reagents should be directed to and will be fulfilled by the Lead Contact, Kenneth Yamada.

### EXPERIMENTAL MODEL AND SUBJECT DETAILS

#### Cell lines and culture

SCC-9 cells were maintained in DMEM/F-12 (Gibco) supplemented with 10% FBS (Hyclone), 2.5 mM L-Glutamine (Gibco), 0.5 mM sodium pyruvate (Gibco) and 400 ng/ml hydrocortisone (Sigma) at 37°C with 5% CO_2_. MDA-MB-231 cells and SW620 cells were maintained in L-15 medium (Gibco) with 10% FBS at 37°C in a 100% air atmosphere (without CO_2_). HT1080 and HFF were cultured in high glucose DMEM (Gibco) with 10% FBS at 37°C with 10% CO_2._ NIH/3T3 cells were cultured in high glucose DMEM with 10% bovine calf serum (Gibco) at 37°C with 10% CO_2._ MCF 10A cells were cultured in the MEGM kit (Lonza) at 37°C with 5% CO_2._ FHC cells were cultured in DMEM/F12 medium supplemented with 10% FBS, additional 10 mM HEPES (to a final concentration of 25 mM), 10 ng/ml cholera toxin (Sigma), 0.005 mg/ml insulin (Thermo Fisher), 0.005 mg/ml transferrin (Thermo Fisher), 100 ng/ml hydrocortisone (Sigma) and 20 ng/ml human recombinant EGF (Thermo Fisher) at 37°C with 5% CO_2_.

Generation of the NIH/3T3 cell line expressing GFP-paxillin was described previously (Doyle et al., 2009). SCC-9 cells expressing GFP-paxillin were generated using the Paxillin-GFP Lentiviral Biosensor (Millipore) according to the manufacturer++’s protocol. Integrin α5 knock-out SCC-9 cell pool and integrin α3 knock-out SCC-9 cell pool were generated under contract by Synthego using CRISPR/Cas9 technology by targeting exon4 of human ITGA5 and exon2 of human ITGA3, respectively, by gRNAs (UCUGUGCGCCAGCUGUACAG and CGGGCACAGCGAGCUCCCGG, respectively). Both myosin IIA and myosin IIB knock-out HFFs were generated by CRISPR-Cas9 targeted genome editing using lentiviral constructs kindly provided by Matthew Kutys at Boston University.

### METHOD DETAILS

#### Basement membrane

Basement membrane was prepared by stripping the overlying mesothelial cells from mouse amniotic membrane using 1N NH_4_OH and mounting the isolated amniotic membrane on glass-bottom MatTek dishes pre-coated with 5 µg/cm^2^ Cell-Tak adhesive (Corning).

#### Extracellular matrix protein coating and patterned coating

Matrigel, laminin, collagen IV, collagen I, perlecan and vitronectin were diluted to 20 µg/ml or other indicated concentrations and aliquoted into glass-bottom MatTek dishes or chambers (ibidi). After incubating at 37°C for 1hr, excess liquid was removed, washed once with PBS, and blocked with 10 µg/ml heat-denatured BSA for 30 mins in room temperature.

For patterned coating, drops of Matrigel were first spotted on glass for 1 hr, which remained as round droplets due to surface tension. After rinsing, vitronectin was added to cover the entire surface for another hour. Since vitronectin does not bind to Matrigel components, this process generates a substrate with patterned coating. Some cells, by chance, adhered and spread on the substrate boundary between Matrigel and vitronectin. A variation of this local substrate patterning was produced by scratching a Matrigel-coated region with a pipette tip followed by incubation with vitronectin to produce a side-by-side Matrigel-vitronectin-Matrigel substrate.

#### Cell invasion assay

5000 SCC-9 cells were seeded in BioCoat Matrigel invasion 24-well chambers (Corning) with 200 µg/ml of the indicated antibodies in serum-free medium; the wells of the plate contained fibronectin-free complete culture medium. After incubating at 37°C for 48 hrs, non-invading cells were removed using cotton swabs. Cells on the lower surface of the membrane were fixed with glutaraldehyde and stained with crystal violet. Cells were photographed at 50X magnification, and all cells within each microscopy field were counted.

#### Fluorescent labeling of fibronectin

Fibronectin purified from human plasma was dialyzed against labeling buffer at pH=8.3 (obtained by mixing 20 parts of PBS with 1 part of 0.2 M sodium bicarbonate solution adjusted to pH=9.0 with sodium hydroxide). To obtain a ratio of labeling of 2-3, a threefold molar excess of atto-565 (Sigma) was added to the fibronectin solution and incubated at room temperature for 12 hours. Unbound dye was removed by gel filtration. The protein solution was then centrifuged at 100,000 rpm in an Airfuge (Beckman Coulter) to remove aggregates. The final concentration and degree of labeling were then calculated and used for experiments.

#### Immunostaining and confocal microscopy

Cells were fixed in 4% paraformaldehyde (Electron Microscopy Sciences) /5% sucrose for 20 mins. Cells were rinsed with Dulbecco’s PBS with calcium and magnesium (PBS++) 3 times and permeabilized with 0.1% Triton X-100 in PBS for 10 mins. Non-specific sites were blocked with 10% donkey serum (Jackson ImmunoResearch Laboratories). Primary and secondary antibodies were diluted in PBS++ with 10% donkey serum for 1 hr and 30 mins, respectively. Cells incubated in MatTek dishes or ibidi chambers were immersed in non-hardening mounting medium (ibidi). Cells on coverslips were mounted onto slides using ProLong Gold Antifade Mountant (Thermo Fisher). Images were collected using an Nikon A1R-MP confocal microscope with 405, 488, 561 and 642 laser lines and either a 60X oil (NA. 1.4) or a 60X water objective (NA. 1.2). The system was controlled by NIS-Elements software (Nikon). A Nikon N-STORM system was also used and is described below.

#### Live cell imaging

SCC-9 cells and NIH/3T3 cells expressing GFP-paxillin were cultured in phenol red-free medium containing 20 µg/ml fibronectin and OxyFluor (Oxyrase) at 1:100 accompanied by 10 mM DL-Lactate (Sigma) to reduce the formation of oxygen free radicals during live imaging. Atto-565 labeled-fibronectin was added to the medium at 5 µg/ml when needed. To visualize actin dynamics, cells were first labeled with SiR-actin at 1 µM for 1 hour. Cell were washed with PBS++ once and then incubated in medium containing 100 nM SiR-actin for the entire live-imaging process. Live imaging was performed by either spinning disk confocal microscope or a Nikon N-STORM TIRF microscope. The spinning disk consisted of a Yokogawa CSU-22 scan head (CSU-21: modified by Spectral Applied Research, Inc.) on an automated Olympus IX-81 microscope using a 60X SAPO-Chromat silicone oil objective (N.A. 1.3) equipped with a custom laser launch with 442, 488, 514, 568 and 642 laser lines (built by A.D.D.), a back-thinned EM-CCD camera (Photometrics), a motorized Z-piezo stage (ASI Imaging, Inc., Eugene, OR) and an environmental chamber maintained cells at constant 37°C with 5-10% CO2 and approximately 50% humidity (Precision Plastics, Beltsville MD). All components were controlled by MetaMorph (Molecular Devices, Downington, PA). The Nikon N-STORM microscope consisted of a Nikon Ti-2 microscope frame, a laser launch with 405, 488, 561 and 642 lines, either 60X or 100X oil (NA 1.49) TIRF objectives, and a CMOS (Flash 4 v3, Hamamatsu) or a back-thinned EM-CCD camera (Photometrics). Temperature, humidity and CO_2_ were maintained constant using a Tokai stage-top incubator. The system was controlled by NIS-Elements software (Nikon).

#### Traction force microscopy

Traction force microscopy was performed as previously described (Plotnikov et al., 2012). Briefly, cells were plated on polyacrylamide gels coated with different extracellular matrix proteins. The gel had a stiffness of 4.3 kPa and was embedded with 655 fluorescent microspheres (Life Technologies). Images were acquired by spinning disk confocal microscopy before and after solubilizing the cells with 10% SDS plus 5% Triton X-100. Images of deformed and released substrates were aligned using ImageJ, and movement of the fluorescent microsphere was quantified with the plugin developed by Tseng *et al*. (Tseng et al., 2011).

#### Quantification and statistical analysis

Images were processed and analyzed by Fiji software. To quantify the amount of fibronectin matrix, images were first subjected to Gaussian blur and background subtraction before total fluorescence intensity was measured. The number of fibronectin matrix fibers as well as the average size of fibers were measured using the FIJI built-in particle analysis function “Analyze Particles…”.

Prism 8 by GraphPad software was used for all analyses. Two-tailed *t* tests were used for pairs of conditions and one-way analysis of variance (ANOVA) followed by the Bonferroni post hoc test was used for comparisons between more than two data sets. All error bars indicate SEM. P<0.05 was considered statistically significant. *n* = the total number of data points per experimental condition.

#### Supplemental video legends

**Video S1**

Time-lapse video of an SCC-9 cell transfected with GFP-paxillin (green) on Matrigel showing fibronectin fibrillogenesis by sliding focal adhesions. Culture medium contained Atto-565-labeled fibronectin (5 μg/ml, red).

**Videos S2 and S3**

Time-lapse videos of NIH/3T3 cell transfected with GFP-paxillin (green) seeded on either vitronectin overnight (Video S2) or Matrigel for 2hrs (Video S3) showing the dynamics of focal adhesions and its relationship to fibronectin fibrillogenesis on Matrigel. Culture medium contained Atto-565-labeled fibronectin (5 μg/ml, red).

**Videos S4 and S5**

Time-lapse videos of an SCC-9 cell transfected with GFP-paxillin (green) on Matrigel showing fibronectin fibrillogenesis mediated by a contractile actomyosin winch. Actin stress fibers were labeled with SiR-actin (red). The medium contained Atto-565-labeled fibronectin (5 μg/ml, blue). Scale bars: 10 μm.

**Video S6**

Time-lapse video of the edge of an SCC-9 cell transfected with GFP-paxillin (green) on Matrigel showing fibronectin fibrillogenesis by sliding focal adhesions during cell retraction. The medium contained Atto-565-labeled fibronectin (5 μg/ml, red).

**Figure S1.**
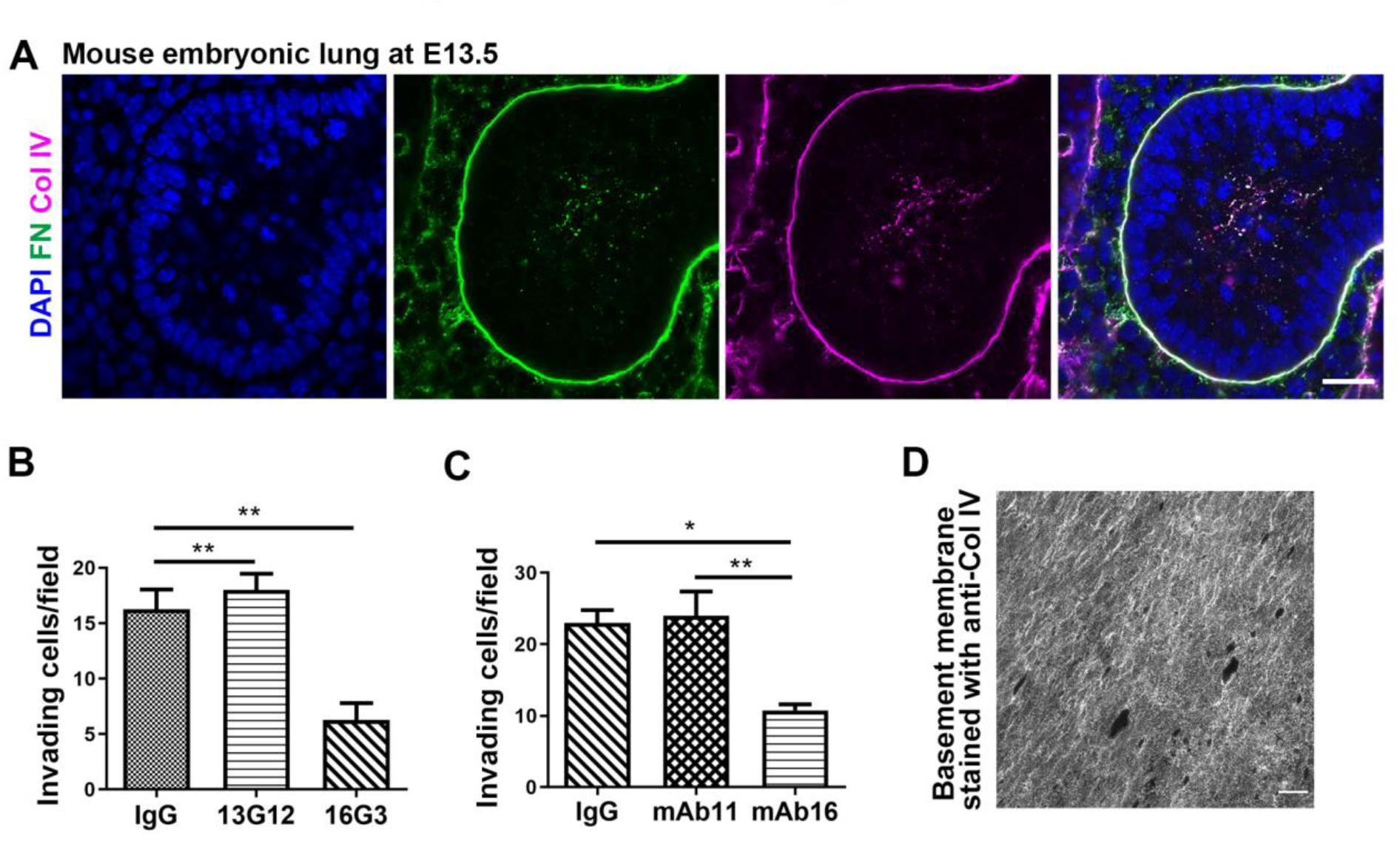
Fibronectin distribution in vivo and a contribution to cancer cell Matrigel invasion. (A) Maximum-projection of confocal images illustrating FN distribution along basement membrane in mouse embryonic lung at stage E13.5. (B) Effects of FN blocking on SCC-9 cell invasion through Matrigel in transwell assay. 13G12: non-functional anti-FN, 16G3: FN blocking antibody. *n*=16. (C) Effects of integrin α5β1 blocking on SCC-9 cell invasion through Matrigel in transwell assays. mAb11: non-functional anti-integrin α5β1, mAb16: integrin α5β1 blocking antibody. Data in (B) and (C) represent average numbers of cells invading across Matrigel transwell per 50x field. *n*=16. (D) Mouse amniotic basement membrane mounted on a tissue culture dish. Error bars: SEM. *p ≤ 0.05, **p ≤ 0.01. Scale bars: 20 μm (A); 40 μm (D).

**Figure S2.**
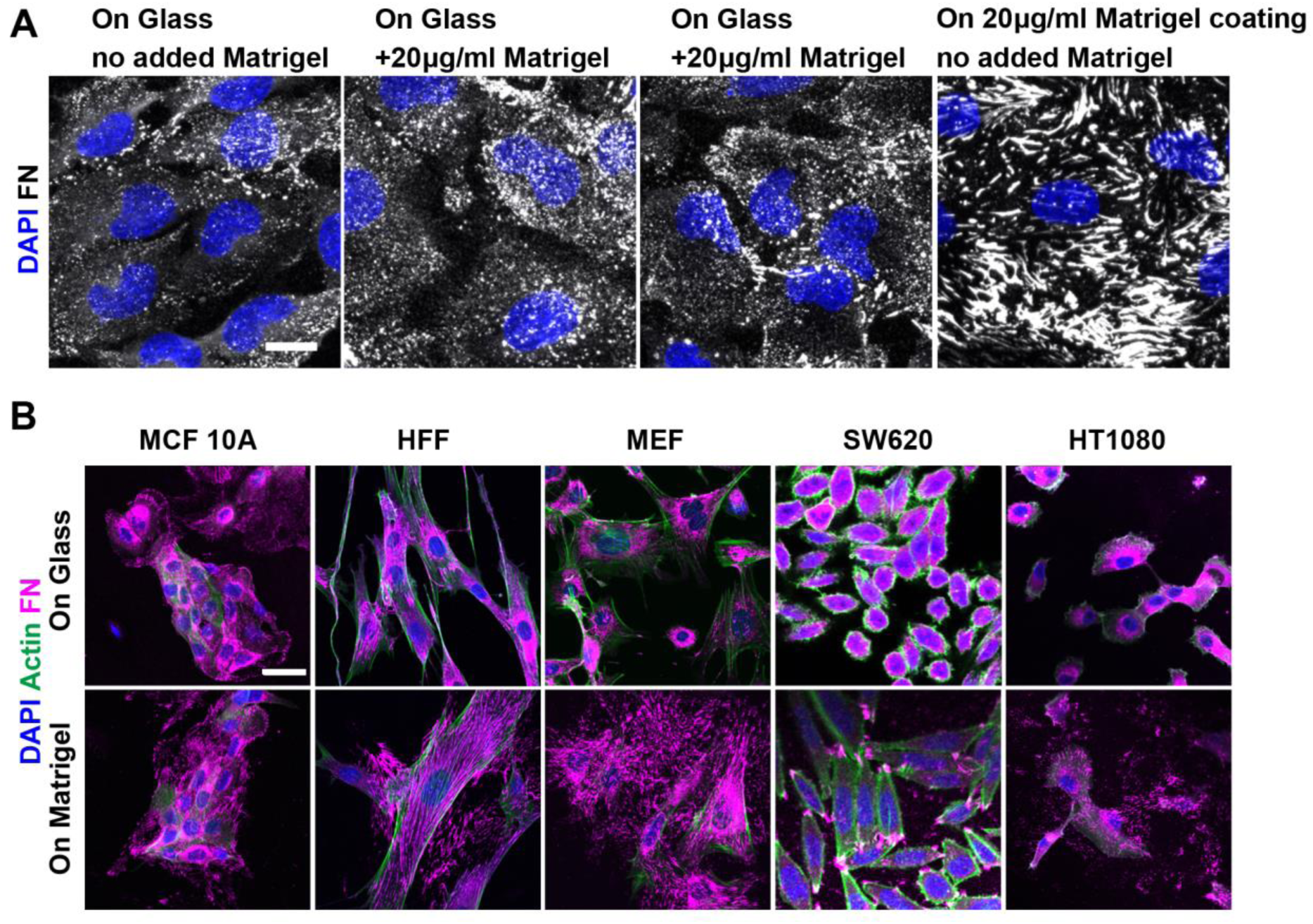
Matrigel as a substrate can promote fibronectin fibrillar matrix deposition. (A) FN fibrillar matrix is induced by Matrigel as a substrate, but not when presented to cells soluble in medium. (B) Effects on FN matrix induction by Matrigel in multiple cell lines. MCF10A: normal human mammary epithelial cells, HFF: human foreskin fibroblasts, MEF: mouse embryonic fibroblasts (E13.5), SW620: human colon cancer cells, HT1080: human fibrosarcoma cells. Compare intracellular FN around the nucleus that is prominent on glass (top row) to matrix deposition on a Matrigel substrate (bottom row). Scale bars: 20 μm (A, B).

**Figure S3.**
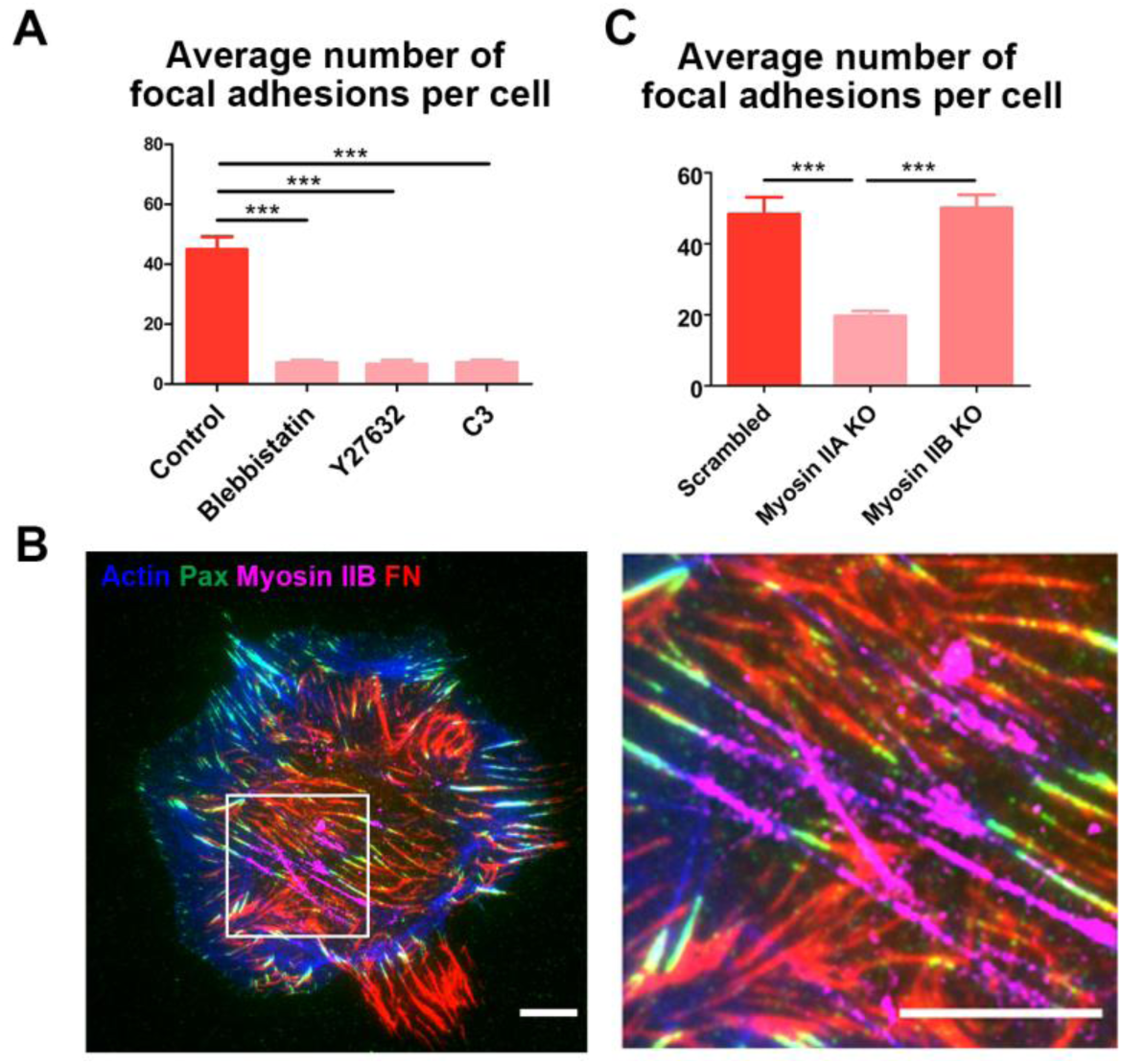
Actomyosin contractility, sliding focal adhesions, and fibronectin matrix assembly on Matrigel. (A) Effects of inhibiting myosin II, Rho-associated protein kinase (ROCK), and Rho by blebbistatin (50 μM), Y27632 (20 μM), or C3 transferase (4μg/ml) on the number of focal adhesions on Matrigel. *n*≥30. (B) TIRF images showing immunolocalization of limits amounts of myosin IIB in association with actin stress fibers positioned between focal adhesions on Matrigel. (C) Effects on myosin IIA and IIB knock-out, respectively, on the number of focal adhesions on Matrigel. *n*=15. Error bars: SEM. ***p ≤ 0.001. Scale bars: 10 μm.

**Figure S4.**
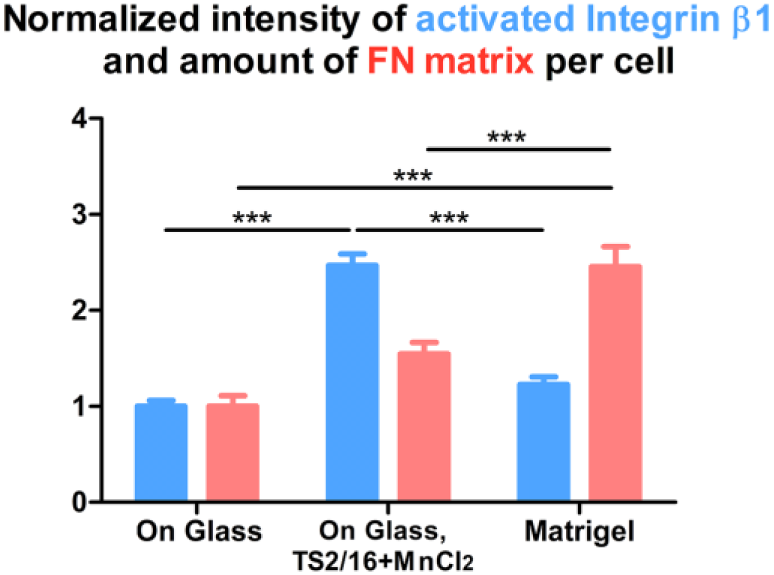
Induction of fibronectin matrix assembly by Matrigel is not due to increased β1 integrin activity. Quantification of β1 integrin activity (by measuring fluorescence intensity of anti-activated β1 integrin-9EG7 staining (blue) and FN matrix assembly (red) by SCC-9 cells growing on glass, on glass with cells treated by MnCl_2_ (1mM) and monoclonal antibody TS2/16 (20 μg/ml) in the medium to stimulate β1 integrin activity versus on Matrigel. Error bars: SEM. ***p ≤ 0.001. *n*≥10.

## References

Akiyama, S.K., Yamada, S.S., Chen, W.T., and Yamada, K.M. (1989). Analysis of fibronectin receptor function with monoclonal antibodies: roles in cell adhesion, migration, matrix assembly, and cytoskeletal organization. J Cell Biol 109, 863–875.

Alberts, B., Johnson, A., Lewis, J., Raff, M., Morgan, D., Roberts, K., and Walter, P. (2015). Molecular Biology of the Cell. New York: Garland Science. 1464.

Aplin, J.D., and Campbell, S. (1985). An immunofluorescence study of extracellular matrix associated with cytotrophoblast of the chorion laeve. Placenta 6, 469–479.

Carter, W.G., Wayner, E.A., Bouchard, T.S., and Kaur, P. (1990). The role of integrins alpha 2 beta 1 and alpha 3 beta 1 in cell-cell and cell-substrate adhesion of human epidermal cells. J Cell Biol 110, 1387–1404.

Couchman, J.R., Gibson, W.T., Thom, D., Weaver, A.C., Rees, D.A., and Parish, W.E. (1979). Fibronectin distribution in epithelial and associated tissues of the rat. Arch Dermatol Res 266, 295–310.

Darribere, T., Riou, J.F., Shi, D.L., Delarue, M., and Boucaut, J.C. (1986). Synthesis and distribution of laminin-related polypeptides in early amphibian embryos. Cell Tissue Res 246, 45–51.

Domogatskaya, A., Rodin, S., and Tryggvason, K. (2012). Functional diversity of laminins. Annu Rev Cell Dev Biol 28, 523–553.

Doyle, A.D., Wang, F.W., Matsumoto, K., and Yamada, K.M. (2009). One-dimensional topography underlies three-dimensional fibrillar cell migration. J Cell Biol 184, 481–490.

Duband, J.L., and Thiery, J.P. (1982a). Appearance and distribution of fibronectin during chick embryo gastrulation and neurulation. Dev Biol 94, 337–350.

Duband, J.L., and Thiery, J.P. (1982b). Distribution of fibronectin in the early phase of avian cephalic neural crest cell migration. Dev Biol 93, 308–323.

Fidler, A.L., Boudko, S.P., Rokas, A., and Hudson, B.G. (2018). The triple helix of collagens - an ancient protein structure that enabled animal multicellularity and tissue evolution. J Cell Sci 131.

Fogerty, F.J., Akiyama, S.K., Yamada, K.M., and Mosher, D.F. (1990). Inhibition of binding of fibronectin to matrix assembly sites by anti-integrin (alpha 5 beta 1) antibodies. J Cell Biol 111, 699–708.

Gardiner, R.A., Seymour, G.J., Lavin, M.F., Strutton, G.M., Gemmell, E., and Hazan, G. (1986). Immunohistochemical analysis of the human bladder. Br J Urol 58, 19–25.

Geiger, B., Bershadsky, A., Pankov, R., and Yamada, K.M. (2001). Transmembrane crosstalk between the extracellular matrix--cytoskeleton crosstalk. Nat Rev Mol Cell Biol 2, 793–805.

Geiger, B., and Yamada, K.M. (2011). Molecular architecture and function of matrix adhesions. Cold Spring Harb Perspect Biol 3.

Georgiadou, M., Lilja, J., Jacquemet, G., Guzman, C., Rafaeva, M., Alibert, C., Yan, Y., Sahgal, P., Lerche, M., Manneville, J.B., et al. (2017). AMPK negatively regulates tensin-dependent integrin activity. J Cell Biol 216, 1107–1121.

Giancotti, F.G., and Ruoslahti, E. (1990). Elevated levels of the alpha 5 beta 1 fibronectin receptor suppress the transformed phenotype of Chinese hamster ovary cells. Cell 60, 849–859.

Gil, J., and Martinez-Hernandez, A. (1984). The connective tissue of the rat lung: electron immunohistochemical studies. J Histochem Cytochem 32, 230–238.

Gonzalez, A.M., Bhattacharya, R., deHart, G.W., and Jones, J.C. (2010). Transdominant regulation of integrin function: mechanisms of crosstalk. Cell Signal 22, 578–583.

Gonzalez, A.M., Gonzales, M., Herron, G.S., Nagavarapu, U., Hopkinson, S.B., Tsuruta, D., and Jones, J.C. (2002). Complex interactions between the laminin alpha 4 subunit and integrins regulate endothelial cell behavior in vitro and angiogenesis in vivo. Proc Natl Acad Sci U S A 99, 16075–16080.

Halliday, N.L., and Tomasek, J.J. (1995). Mechanical properties of the extracellular matrix influence fibronectin fibril assembly in vitro. Exp Cell Res 217, 109–117.

Hay, E.D. (1991). Cell Biology of Extracellular Matrix. 2nd ed. Plenum Press, New York. 468.

Horton, E.R., Humphries, J.D., James, J., Jones, M.C., Askari, J.A., and Humphries, M.J. (2016). The integrin adhesome network at a glance. J Cell Sci 129, 4159–4163.

Houdijk, W.P., de Groot, P.G., Nievelstein, P.F., Sakariassen, K.S., and Sixma, J.J. (1986). Subendothelial proteins and platelet adhesion. von Willebrand factor and fibronectin, not thrombospondin, are involved in platelet adhesion to extracellular matrix of human vascular endothelial cells. Arteriosclerosis 6, 24–33.

Houdijk, W.P., and Sixma, J.J. (1985). Fibronectin in artery subendothelium is important for platelet adhesion. Blood 65, 598–604.

Hughes, P.E., Diaz-Gonzalez, F., Leong, L., Wu, C., McDonald, J.A., Shattil, S.J., and Ginsberg, M.H. (1996). Breaking the integrin hinge. A defined structural constraint regulates integrin signaling. J Biol Chem 271, 6571–6574.

Hynes, R.O. (1990). Fibronectins. New York: Springer-Verlag. 544.

Hynes, R.O. (1999). The dynamic dialogue between cells and matrices: implications of fibronectin’s elasticity. Proc Natl Acad Sci U S A 96, 2588–2590.

Ilic, D., Kovacic, B., Johkura, K., Schlaepfer, D.D., Tomasevic, N., Han, Q., Kim, J.B., Howerton, K., Baumbusch, C., Ogiwara, N., et al. (2004). FAK promotes organization of fibronectin matrix and fibrillar adhesions. J Cell Sci 117, 177–187.

Jayadev, R., and Sherwood, D.R. (2017). Basement membranes. Curr Biol 27, R207–R211.

Katz, B.Z., Zamir, E., Bershadsky, A., Kam, Z., Yamada, K.M., and Geiger, B. (2000). Physical state of the extracellular matrix regulates the structure and molecular composition of cell-matrix adhesions. Mol Biol Cell 11, 1047–1060.

Kern, A., Eble, J., Golbik, R., and Kuhn, K. (1993). Interaction of type IV collagen with the isolated integrins alpha 1 beta 1 and alpha 2 beta 1. Eur J Biochem 215, 151–159.

Kleinman, H.K., and Martin, G.R. (2005). Matrigel: basement membrane matrix with biological activity. Semin Cancer Biol 15, 378–386.

Kohno, T., Sorgente, N., Ishibashi, T., Goodnight, R., and Ryan, S.J. (1987). Immunofluorescent studies of fibronectin and laminin in the human eye. Invest Ophthalmol Vis Sci 28, 506–514.

Krotoski, D.M., Domingo, C., and Bronner-Fraser, M. (1986). Distribution of a putative cell surface receptor for fibronectin and laminin in the avian embryo. J Cell Biol 103, 1061–1071.

Languino, L.R., Gehlsen, K.R., Wayner, E., Carter, W.G., Engvall, E., and Ruoslahti, E. (1989). Endothelial cells use alpha 2 beta 1 integrin as a laminin receptor. J Cell Biol 109, 2455–2462.

Leppi, T.J., Repesh, L.A., Furcht, L.T., Bartzen, P.J., and Holt, G.E. (1982). Immunohistochemical localization of fibronectin in human and rat uterine cervices. J Histochem Cytochem 30, 413–417.

Linder, E., Stenman, S., Lehto, V.P., and Vaheri, A. (1978). Distribution of fibronectin in human tissues and relationship to other connective tissue components. Ann N Y Acad Sci 312, 151–159.

Ly, D.P., Zazzali, K.M., and Corbett, S.A. (2003). De novo expression of the integrin alpha5beta1 regulates alphavbeta3-mediated adhesion and migration on fibrinogen. J Biol Chem 278, 21878–21885.

Manabe, R., Tsutsui, K., Yamada, T., Kimura, M., Nakano, I., Shimono, C., Sanzen, N., Furutani, Y., Fukuda, T., Oguri, Y., et al. (2008). Transcriptome-based systematic identification of extracellular matrix proteins. Proc Natl Acad Sci U S A 105, 12849–12854.

Mayer, B.W., Jr., Hay, E.D., and Hynes, R.O. (1981). Immunocytochemical localization of fibronectin in embryonic chick trunk and area vasculosa. Dev Biol 82, 267–286.

McKeown-Longo, P.J., and Mosher, D.F. (1983). Binding of plasma fibronectin to cell layers of human skin fibroblasts. J Cell Biol 97, 466–472.

Meier, S., and Drake, C. (1984). SEM localization of cell-surface-associated fibronectin in the cranium of chick embryos utilizing immunolatex microspheres. J Embryol Exp Morphol 80, 175–195.

Meng, X.N., Jin, Y., Yu, Y., Bai, J., Liu, G.Y., Zhu, J., Zhao, Y.Z., Wang, Z., Chen, F., Lee, K.Y., et al. (2009). Characterisation of fibronectin-mediated FAK signalling pathways in lung cancer cell migration and invasion. Br J Cancer 101, 327–334.

Miles, A.J., Knutson, J.R., Skubitz, A.P., Furcht, L.T., McCarthy, J.B., and Fields, G.B. (1995). A peptide model of basement membrane collagen alpha 1 (IV) 531-543 binds the alpha 3 beta 1 integrin. J Biol Chem 270, 29047–29050.

Mosher, D.F. (1989). Fibronectin. Elsevier. 492.

Nagai, T., Yamakawa, N., Aota, S., Yamada, S.S., Akiyama, S.K., Olden, K., and Yamada, K.M. (1991). Monoclonal antibody characterization of two distant sites required for function of the central cell-binding domain of fibronectin in cell adhesion, cell migration, and matrix assembly. J Cell Biol 114, 1295–1305.

Newgreen, D., and Thiery, J.P. (1980). Fibronectin in early avian embryos: synthesis and distribution along the migration pathways of neural crest cells. Cell Tissue Res 211, 269–291.

Ohashi, T., Kiehart, D.P., and Erickson, H.P. (2002). Dual labeling of the fibronectin matrix and actin cytoskeleton with green fluorescent protein variants. J Cell Sci 115, 1221–1229.

Pankov, R., Cukierman, E., Katz, B.Z., Matsumoto, K., Lin, D.C., Lin, S., Hahn, C., and Yamada, K.M. (2000). Integrin dynamics and matrix assembly: tensin-dependent translocation of alpha(5)beta(1) integrins promotes early fibronectin fibrillogenesis. J Cell Biol 148, 1075–1090.

Pankov, R., and Yamada, K.M. (2002). Fibronectin at a glance. J Cell Sci 115, 3861–3863.

Plotnikov, S.V., Pasapera, A.M., Sabass, B., and Waterman, C.M. (2012). Force fluctuations within focal adhesions mediate ECM-rigidity sensing to guide directed cell migration. Cell 151, 1513–1527.

Pozzi, A., Yurchenco, P.D., and Iozzo, R.V. (2017). The nature and biology of basement membranes. Matrix Biol 57-58, 1–11.

Schwarzbauer, J.E., and Sechler, J.L. (1999). Fibronectin fibrillogenesis: a paradigm for extracellular matrix assembly. Curr Opin Cell Biol 11, 622–627.

Sechler, J.L., Corbett, S.A., and Schwarzbauer, J.E. (1997). Modulatory roles for integrin activation and the synergy site of fibronectin during matrix assembly. Mol Biol Cell 8, 2563–2573.

Sechler, J.L., Cumiskey, A.M., Gazzola, D.M., and Schwarzbauer, J.E. (2000). A novel RGD-independent fibronectin assembly pathway initiated by alpha4beta1 integrin binding to the alternatively spliced V region. J Cell Sci 113 (Pt 8), 1491–1498.

Singh, P., Carraher, C., and Schwarzbauer, J.E. (2010). Assembly of fibronectin extracellular matrix. Annu Rev Cell Dev Biol 26, 397–419.

Smilenov, L.B., Mikhailov, A., Pelham, R.J., Marcantonio, E.E., and Gundersen, G.G. (1999). Focal adhesion motility revealed in stationary fibroblasts. Science 286, 1172–1174.

Staatz, W.D., Walsh, J.J., Pexton, T., and Santoro, S.A. (1990). The alpha 2 beta 1 integrin cell surface collagen receptor binds to the alpha 1 (I)-CB3 peptide of collagen. J Biol Chem 265, 4778–4781.

Takada, Y., Murphy, E., Pil, P., Chen, C., Ginsberg, M.H., and Hemler, M.E. (1991). Molecular cloning and expression of the cDNA for alpha 3 subunit of human alpha 3 beta 1 (VLA-3), an integrin receptor for fibronectin, laminin, and collagen. J Cell Biol 115, 257–266.

Tseng, Q., Wang, I., Duchemin-Pelletier, E., Azioune, A., Carpi, N., Gao, J., Filhol, O., Piel, M., Thery, M., and Balland, M. (2011). A new micropatterning method of soft substrates reveals that different tumorigenic signals can promote or reduce cell contraction levels. Lab Chip 11, 2231–2240.

Uscanga, L., Kennedy, R.H., Stocker, S., Grimaud, J.A., and Sarles, H. (1984). Immunolocalization of collagen types, laminin and fibronectin in the normal human pancreas. Digestion 30, 158–164.

Warburton, M.J., Ormerod, E.J., Monaghan, P., Ferns, S., and Rudland, P.S. (1981). Characterization of a myoepithelial cell line derived from a neonatal rat mammary gland. J Cell Biol 91, 827–836.

Wartiovaara, J., Leivo, I., Virtanen, I., Vaheri, A., and Graham, C.F. (1978). Appearance of fibronectin during differentiation of mouse teratocarcinoma in vitro. Nature 272, 355–356.

Wennerberg, K., Lohikangas, L., Gullberg, D., Pfaff, M., Johansson, S., and Fassler, R. (1996). Beta 1 integrin-dependent and -independent polymerization of fibronectin. J Cell Biol 132, 227–238.

Wierzbicka-Patynowski, I., Mao, Y., and Schwarzbauer, J.E. (2007). Continuous requirement for pp60-Src and phospho-paxillin during fibronectin matrix assembly by transformed cells. J Cell Physiol 210, 750–756.

Wierzbicka-Patynowski, I., and Schwarzbauer, J.E. (2002). Regulatory role for SRC and phosphatidylinositol 3-kinase in initiation of fibronectin matrix assembly. J Biol Chem 277, 19703–19708.

Winograd-Katz, S.E., Fassler, R., Geiger, B., and Legate, K.R. (2014). The integrin adhesome: from genes and proteins to human disease. Nat Rev Mol Cell Biol 15, 273–288.

Wu, C., Chung, A.E., and McDonald, J.A. (1995a). A novel role for alpha 3 beta 1 integrins in extracellular matrix assembly. J Cell Sci 108 (Pt 6), 2511–2523.

Wu, C., Hughes, P.E., Ginsberg, M.H., and McDonald, J.A. (1996). Identification of a new biological function for the integrin alpha v beta 3: initiation of fibronectin matrix assembly. Cell Adhes Commun 4, 149–158.

Wu, C., Keivens, V.M., O’Toole, T.E., McDonald, J.A., and Ginsberg, M.H. (1995b). Integrin activation and cytoskeletal interaction are essential for the assembly of a fibronectin matrix. Cell 83, 715–724.

Yang, J.T., Rayburn, H., and Hynes, R.O. (1993). Embryonic mesodermal defects in alpha 5 integrin-deficient mice. Development 119, 1093–1105.

Yoneda, A., Ushakov, D., Multhaupt, H.A., and Couchman, J.R. (2007). Fibronectin matrix assembly requires distinct contributions from Rho kinases I and -II. Mol Biol Cell 18, 66–75.

Zaidel-Bar, R., and Geiger, B. (2010). The switchable integrin adhesome. J Cell Sci 123, 1385–1388.

Zaidel-Bar, R., Milo, R., Kam, Z., and Geiger, B. (2007). A paxillin tyrosine phosphorylation switch regulates the assembly and form of cell-matrix adhesions. J Cell Sci 120, 137–148.

Zamir, E., Katz, M., Posen, Y., Erez, N., Yamada, K.M., Katz, B.Z., Lin, S., Lin, D.C., Bershadsky, A., Kam, Z., et al. (2000). Dynamics and segregation of cell-matrix adhesions in cultured fibroblasts. Nat Cell Biol 2, 191–196.

Zhang, Q., Peyruchaud, O., French, K.J., Magnusson, M.K., and Mosher, D.F. (1999). Sphingosine 1-phosphate stimulates fibronectin matrix assembly through a Rho-dependent signal pathway. Blood 93, 2984–2990.

Zhong, C., Chrzanowska-Wodnicka, M., Brown, J., Shaub, A., Belkin, A.M., and Burridge, K. (1998). Rho-mediated contractility exposes a cryptic site in fibronectin and induces fibronectin matrix assembly. J Cell Biol 141, 539–551.

